# PangeBlocks: customized construction of pangenome graphs via maximal blocks

**DOI:** 10.1101/2024.09.17.613426

**Authors:** Jorge Avila Cartes, Paola Bonizzoni, Simone Ciccolella, Gianluca Della Vedova, Luca Denti

**Author notes:** Contributing authors.

## Abstract

**Background:** The construction of a pangenome graph is a fundamental task in pangenomics. A natural theoretical question is how to formalize the computational problem of building an optimal pangenome graph, making explicit the underlying optimization criterion and the set of feasible solutions. Current approaches build a pangenome graph with some heuristics, without assuming some explicit optimization criteria. Thus it is unclear how a specific optimization criterion affects the graph topology and downstream analysis, like read mapping and variant calling.

**Methods:** In this paper, by leveraging the notion of maximal block in a Multiple Sequence Alignment (MSA), we reframe the pangenome graph construction problem as an exact cover problem on blocks called *Minimum Weighted Block Cover* (MWBC). Then we propose an Integer Linear Programming (ILP) formulation for the MWBC problem that allows us to study the most natural objective functions for building a graph.

**Results:** We provide an implementation of the ILP approach for solving the MWBC and we evaluate it on SARS-CoV-2 complete genomes, showing how different objective functions lead to pangenome graphs that have different properties, hinting that the specific downstream task can drive the graph construction phase.

**Conclusion:** We show that a customized construction of a pangenome graph based on selecting objective functions has a direct impact on the resulting graphs. In particular, our formalization of the MWBC problem, based on finding an optimal subset of blocks covering an MSA, paves the way to novel practical approaches to graph representations of an MSA where the user can guide the construction.

## 1 Introduction

The notion of a microbial pangenome goes back to 2005 [1], and it is usually considered the first example of the use of the term pangenome to describe multiple genomes. Under this definition, the microbial pangenome comprises two fundamental components: the core genome, encompassing the genes that are ubiquitous across all strains within a given microbial population, and the dispensable genomes, which encompass genes that are absent in at least one strain within the population.

Nowadays sequencing organisms is essentially routine, as we have witnessed during the SARS-CoV-2 pandemic, when millions of viral genomes have been sequenced. The introduction of Next-Generation Sequencing (NGS) technologies in 2006 made sequencing cheaper and more accessible. Later on, a new sub-area of research in computational biology was consolidated to address the intrinsic challenges introduced by the availability of several genomes, named computational pangenomics. In computational pangenomics, a *pangenome* is a collection of genomic sequences to be analyzed jointly, or to be used as a reference [2]. The natural representation of a pangenome is a directed graph [3], therefore computational pangenomics highlights the transition from working with reference genomes as strings to pangenome graphs.

Pangenome graphs have demonstrated their ability to encompass more comprehensive information, notably in the domain of crops [4], bovine [5] and human [6] data, with important implications for accurate identification of structural variations, especially when contrasted with conventional linear reference genome assemblies. Furthermore, a noteworthy issue associated with linear reference genome assemblies is the lack of genetic diversity that characterizes, for example, the human reference, but is especially evident in bacterial pangenomes. Specifically, it has been empirically established that the current human reference (GRCh38) inadequately captures the genetic diversity within the African population [7].

The need for richer data structures to represent pangenomes is clear. However, the pursuit of establishing an unambiguous notion of *optimal* pangenome graph is still unsettled. For example, pangenome graphs can be built with different tools (`vg` [8], `minigraph` [9], `pggb` [10], `minigraph-cactus` [6], `make_prg` [11], `founderblockgraph` [12]), but the comparison of their results has always relied on specific downstream analyses [5, 6, 13], while we would like to have a quality measure that is not overly dependent on downstream analysis.

It is noteworthy that computational tools diverge in several critical attributes, including scalability, the ability to represent exactly the original genomes, the presence of cycles in the graph, and the proficiency in showcasing well-documented genetic variants. In particular, a critical aspect to consider in the construction of a pangenome graph is whether the graph is suitable for tasks such as mapping reads under the seedand-extend paradigm. In fact, when most of the vertices have a label that is shorter than the seed length, obtaining a high quality alignment becomes harder [8, 14–16].

We propose a formal framework to build a variation graph [3]) that (1) is acyclic and (2) represents perfectly the input genomes, from a Multiple Sequence Alignment (MSA) of the input genomes. More precisely, we frame the pangenome construction problem as finding an optimal tiling of the MSA into blocks, where each block is the set of cells of the MSA defined by a set of rows and an interval of consecutive columns, where all rows of the block are labeled by the same string. Each block becomes a vertex of the pangenome graph and we add arcs to the graph to connect blocks that are adjacent in the MSA. This guarantees that each input sequence corresponds to a path of an acyclic graph. More precisely, we introduce an optimization problem, the Minimum Weight Block Cover problem, in short MWBC, which, given a set of blocks, finds an optimal subset of blocks that cover the MSA. The MWBC problem is a special case of a more general optimization problem, called General Minimum Weight Block Cover problem that asks for a set of blocks that cover the MSA, where the set is optimum w.r.t. a weighted function defined over the solution.

Other approaches, based on Elastic Founder Graphs (EFG) [12, 17, 18] use the notion of block as a portion of the MSA resulting from a vertical segmentation of it. Under this definition, a block is essentially a subMSA instance, where several strings might belong to a block, while our definition of block is based on maximal blocks and relates to a unique string. The EFG approach aims to create indexable graphs for linear time pattern matching queries and propose several optimization criteria for the segmentation of the MSA for enhancing the efficiency and expressiveness of the graph for queries.

Currently, `make_prg` [11] and `founderblockgraph` [12] are the only tools that guarantee the construction of an acyclic graph. The former builds a sequence graph, which is a graph whose paths are not distinguished. On the other hand, the second one creates a variation graph with all sequences explicitly expressed as a path in the graph, but both express all input sequences in the final graph. On the other hand, `vg` can remove cycles from a variation graph, but only at a later step: its main heuristics are tailored for the general case.

The MWBC problem is similar to some problems that have been studied in the literature, such as the Weighted Rectangle Cover [19] or the Maximum Weighted Submatrix Coverage [20] problems. Unfortunately, none of those problems are on matrices containing characters and where the order of the rows is irrelevant, therefore MWBC does not inherit their hardness results. On the other hand, the similarities are sufficiently strong that we conjecture that MWBC is NP-hard.

Besides the formulation of the MWBC problem, we propose an ILP approach for solving MWBC, based on the classical ILP formulation of Exact Set Cover [21]. To avoid including all possible blocks of an MSA in the formulation, first we identify its maximal blocks, a notion introduced in the context of haplotyping, since they can be computed in linear time [22].

More precisely, we use an extension, called Wild-pBWT^1^, that works on an arbitrary alphabet, with this approach we can compute blocks directly over an MSA over the alphabet Σ = {*A, C, G, T, N*} extended with the special symbol − called *indel*. Hence characters *N* and − are treated equally as *ACGT* for computing blocks.

Moreover, we restrict our attention to blocks that are obtained by decomposing overlapping maximal blocks. We will describe two decomposition strategies; a slower strategy that produces more blocks, but is limited to smaller instances, and a faster strategy that can be applied to larger MSAs. The result of the decomposition phase is a set of blocks that is an instance of MWBC-B.

We propose and analyze five different objective functions for the ILP formulation, each tailored for a different goal. Those objective functions aim to highlight some measurable properties that the graph should exhibit, such as the length of the labels, the number of paths traversing the vertices, and the total size of the graph. These are the first elementary properties that can be measured or optimized when constructing the graph. On the other hand, an open question is how can we computationally encode in the graph construction some properties that are biological meaningfully, such as how to represent similarities or differences between the sequences or some biological events.

We assess experimentally the impact on the final pangenome graph of the different strategies used to produce the set of blocks that is an instance of the MWBC problem. Then we measure the impact of the objective functions on an MSA built from 50 complete SARS-CoV-2 genomes — we have obtained similar results on 20 and 100 complete SARS-CoV-2 genomes.

Our results show that a strategy that produces a larger set of blocks improves significantly the optimal values reached by each objective function at the expense of an increase in the computation time for solving the MWBC problem.

Most notably, with `pangeblocks` we were able to construct variation graphs where the number of potential seeds is much larger than the graphs computed by `pggb`— this is a property that helps tools based on seeding approaches like GraphAligner [14]. We also notice that graphs built with `vg` are closer (w.r.t. the properties measured) to the graphs built with `pangeblocks` when minimizing the length of the graph. On the other hand, compared to `pggb`, `pangeblocks` is able to produce graphs with a smaller number of nodes in general, and in particular has significantly fewer nodes that are used by only a smaller percentage of the input genome sequences. In conclusion, the experimental analysis shows that pangenome graph measures may significantly change with the use of objective functions in building an optimal graph under such functions.

## 2 Preliminaries

In what follows, we recall and give some definitions needed to define the problem we are solving. We start with the definitions of variation graph and sequence graph [3]. In the paper, we deal with strings over the alphabet Σ = {*A, C, G, T, N* } extended with the special symbol − called *indel* that is not in Σ. The *length* of the string *s*, denoted by |*s*|, is the number of characters of *s*.

Given a string s, we denote *s*[*b* : *e*] the substring of *s* from position *b* to position *e*, where 1 ≤ *b* ≤*e* ≤ |*s*|, and the *i*-esim element of *s* is denoted by *s*[*i*].

### Definition 1

(Variation Graph [3]). *A variation graph G* = ⟨*V, A, W* ⟩*is a directed graph whose vertices are labeled by nonempty strings, with λ* : *V*↦ Σ^+^ *being the labeling function, and where A denotes the set of arcs and W denotes a nonempty set of distinguished walks*.

A *sequence graph G* = ⟨*V, A*⟩ is a graph obtained from a variation graph by ignoring the set *W* of distinguished walks. Let *G* be a variation graph and let *w* = ⟨*v*_1_, …, *v*_*l*_⟩ be a walk of *G*. Then the *label* of the walk *w* is the concatenation *λ*(*w*) = *λ*(*v*_1_) · · · *λ*(*v*_*l*_) of the labels of the vertices of the walk. Given the string *g*, then *G expresses g* if there is a walk *w* ∈ *W* such that the label of the walk *w* is exactly *g*, that is *λ*(*w*) = *g*.

Since our algorithm starts from an MSA, we need to provide a formal definition of *multiple sequence alignment*, while referring the reader to [24] for a more detailed exposition.

### Definition 2

(Expansion of a sequence). *Given a string s* = *s*[1]*s*[2] · · · *s*[*n*] *over the alphabet* Σ, *an* expansion *t of s is obtained by possibly inserting indels into s. A maximal substring of t containing only indels is called a gap*.

### Definition 3

(Multiple Sequence Alignment (MSA)). *Let* = 𝒮 {*s*_1_, …, *s*_*m*_} *be a set of k strings. Then an MSA* 𝒮 *of is a set* 𝒯 = {*t*_1_, …, *t*_*m*_} *such that each t*_*i*_ *is an expansion of s*_*i*_, *all t*_*i*_ *have the same length n (also called the* length *of the MSA) and for all j with* 1 ≤ *j* ≤ *n, then there exists a t*_*i*_ *such that t*_*i*_[*j*] ≠ −.

We will use the example of Figure 1 as a running example to introduce the technical aspects of this paper.

**Fig. 1:**
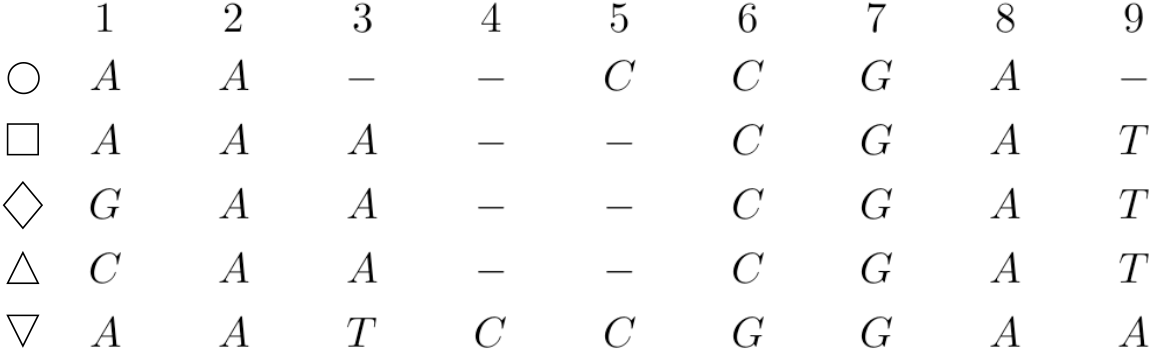
Example of MSA on the sequences ∘= *AACCGA*, □ = *AAACGAT*, ♦ = *GAACGAT*, △ = *CAACGAT*, and ▽ = *AATCCGGAA*.

**Fig. 2:**
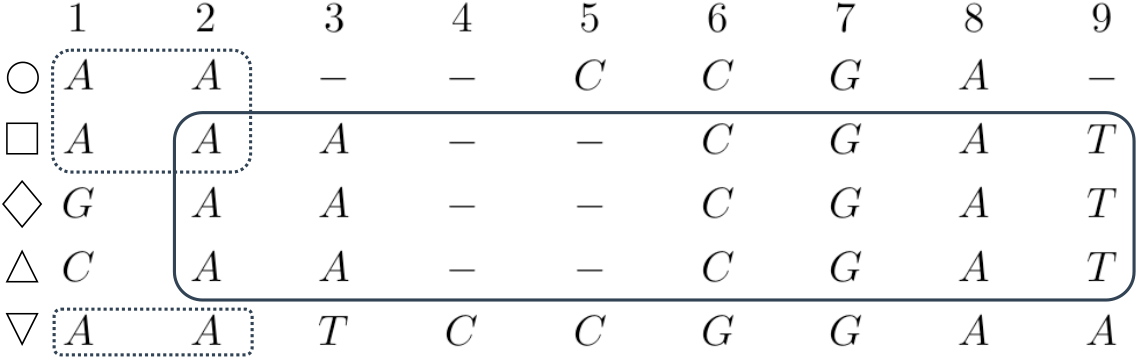
Two overlapping maximal blocks whose sets of rows are not nested. On the left, the (dashed line) block ({∘, □, ▽}, 1, 2), on the right the (solid line) block ({□, ♦, △}, 2, 9)

**Fig. 3:**
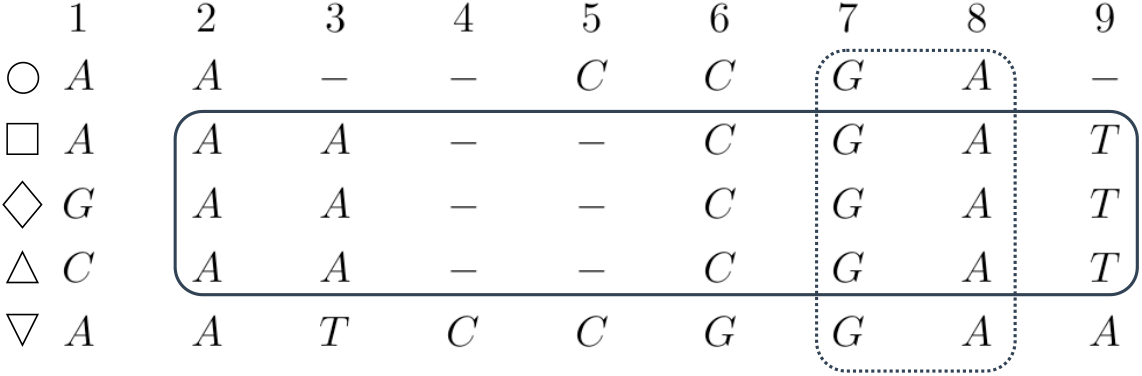
Two overlapping maximal blocks with nested sets of rows Starting from the left, we have the red block ({ □, ♦,△},, 2, 9) and the blue block ({∘, □, ♦, △, ▽ }, 7, 8). The blue block is a *vertical block* because it contains all rows of the MSA.

A variation graph is a representation of an MSA [25], but such a representation is not unique. Indeed there can exist more than one variation graph expressing the sequences in an MSA. Conversely, from each variation graph we can obtain several MSAs, *e*.*g*. depending on where indels are inserted. However, it is not immediate to obtain those alignments, since a graph might contain cycles that must be broken to obtain the MSA.

In this paper we explore a connection between MSAs and variation graphs based on the notion of *block*, where a block is a portion of the MSA correponding to the same string and is associated with a vertex of the variation graph.

In what follows, given a set 𝒮 = {*s*_1_, …, *s*_*k*_} of sequences and a subset *K* ⊂ {1, …, *k* }of indexes, 𝒮 | _*K*_ is defined as the set {*s*_*i*_ : *I* ∈ *K*}. We can now introduce the definition of block.

### Definition 4

(Block). *Let* 𝒯= {*t*_1_, …, *t*_*m*_}*be an MSA of length n. Then a block is a triple* (*K, b, e*) *with K* ⊆ {1, …, *m*}, *K* ≠ ∅*and* 1 ≤ *b* ≤ *e* ≤ *n, such that t*_*i*_[*b* : *e*] = *t*_*j*_[*b* : *e*] *for all i, j* ∈ *K*.

The *label* of the block (*K, b, e*) is the string *t*_*l*_[*b* : *e*], for any *l*∈ *K*.

Informally, a block is a collection *K* of rows of the MSA and an interval of columns between *b* and *e* such that all rows of *K* in that interval are the same sequence.

In this paper, each vertex of the variation graph corresponds to a block, and the label of the vertex is exactly the label of the block. Moreover, the vertices of the graph correspond to non-overlapping blocks of the MSA. While the number of blocks is exponential in the number of sequences, maximal blocks can be computed in linear time, where a block (*K, b, e*) is maximal if enlarging *K*, decreasing *b*, or increasing *e* does not result in a block.

### Definition 5

(Maximal block). *Let* 𝒯= {*t*_1_, …, *t*_*m*_} *be an MSA of length n, and let* (*K, b, e*) *be a block of* 𝒯. *Then* (*K, b, e*) *is a maximal block if:*

1. *for each h* ∈ {1, …, *m*} \ *K, t*_*h*_[*b* : *e*] ≠ *t*_*k*_[*b* : *e*] *for any k* ∈ *K (row-maximality)*,
2. *b* = 1 *or t*_*i*_[*b* − 1] ≠ *t*_*j*_[*b* − 1] *for some t*_*i*_, *t*_*j*_ ∈ 𝒯 _*K*_ *(left-maximality), and*
3. *e* = *n or t*_*i*_[*e* + 1] ≠ *t*_*j*_[*e* + 1] *for some t*_*i*_, *t*_*j*_ ∈ 𝒯_*K*_ *(right-maximality)*.

Finally, we say that two blocks are overlapping if they share some entry of the MSA.

### Definition 6

(Overlapping blocks). *Two blocks* (*K*_1_, *b*_1_, *e*_1_) *and* (*K*_2_, *b*_2_, *e*_2_) overlap *if*

*(1) K*_1_ ∩*K*_2_ ≠ ∅ *and (2) there exists an integer i such that b*_1_ ≤ *i* ≤ *e*_1_ *and b*_2_ ≤ *i* ≤ *e*_2_.

## 3 The MWBC problem

We formalize the variation graph construction as computing a block cover of the MSA. **Definition 7** (Block Cover). *Let* 𝒯 = {*t*_1_, …, *t*_*m*_} *be an MSA of length n over m sequences, and let* B *be a set of blocks of* 𝒯. *Then ℬ is a* cover *of the MSA* 𝒯 *if, for each* 1 ≤ *i* ≤ *m and* 1 ≤ *j* ≤ *n, there exists at least a block* (*K, b, e*) *such that i* ∈ *K and b* ≤ *j* ≤ *e. Moreover, if all blocks in* ℬ *are non-overlapping, then* ℬ *is an* exact cover *of the MSA*.

The natural computational problem that we formalize is the Minimum Weight Block Cover, where the instance is an MSA and we want to compute an exact cover of the MSA using blocks that minimize some objective function.

Notice that we are not specifying the objective function here, since different biological settings will result in different objecting functions. In fact, one of the advantages of our approach is that we can easily consider different objective functions, resulting in the following (meta)problem.

### Problem 1

(General Minimum Weight Block Cover (G-MWBC)). *The instance of the G-MWBC problem is an MSA* 𝒯. *Then a feasible solution is an exact cover 𝒞 of* 𝒯. *We denote f* () *as the objective function. Then 𝒞 is an optimal solution if 𝒞 minimizes the objective function*.

Our strategy to solve G-MWBC will mainly consist of selecting a set of block ℬ and looking for a subset of that is an exact cover of ℬ the MSA. This is formalized with the following computational problem, where the set of possible blocks is given as part of the instance.

### Problem 2

(Minimum Weight Block Cover given a set of blocks (MWBC)). *The instance of the MWBC problem is an MSA* 𝒯 *and a set* ℬ*of blocks of the MSA. Then a feasible solution, if it exists, is an exact cover* 𝒞⊆ *ℬof* 𝒯. *We denote f* () *as the objective function. Then is an optimal solution if* 𝒞 *minimizes the objective function*.

As already observed, we conjecture that G-MWBC and MWBC are NP-hard even though they are restricted versions of the (weighted) exact set cover. In fact, we can easily encode an instance of MWBC as an instance of minimum weight exact set cover, and G-MWBC is the special case of MWBC where ℬ is the set of all possible blocks of the MSA.

### 3.1 Building a variation Graph from a Block Cover

Once we have an exact cover 𝒞 of the MSA (*e*.*g*. Figure 4), we compute a variation graph whose vertices are exactly the blocks of the cover, and two blocks (*K*_1_, *b*_1_, *e*_1_) and (*K*_2_, *b*_2_, *e*_2_) are an arc of the graph iff *e*_1_ = *b*_2_ − 1, and *K*_1_ ∩ *K*_2_ ≠ ∅, that is those blocks correspond to consecutive columns of the MSA, and share at least one row. Observe that the variation graph we build is a directed acyclic graph.

**Fig. 4:**
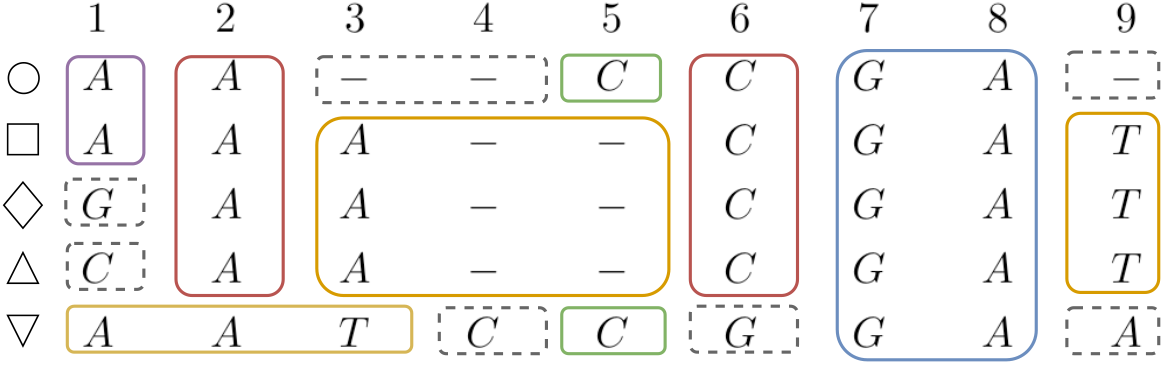
Exact cover of the MSA of Figure 1. Blocks with a solid line contain at least 2 rows. Blocks with dashed lines contain only 1 row — they are one-row or one-character blocks. The same colors will be used for the associated variation graph (see Figures 5 and 6).

The label of a vertex is the label of the corresponding block, as shown in Figure 5. Observe that, by construction, each row of the MSA is the concatenation of the labels of blocks corresponding to a path of the graph. This means that the graph we have obtained is a lossless representation of the input sequences of the MSA. Although the graph represents each input sequence without any loss of information, it might require some post-processing steps to obtain a simpler graph that is able to represent the same set of sequences. First, we remove all nodes whose label consists of indels only — when such a node is removed, an arc is added between its predecessors and successors, to preserve the property that the graph expresses all the input sequences. Then we remove indels from all labels.

**Fig. 5:**
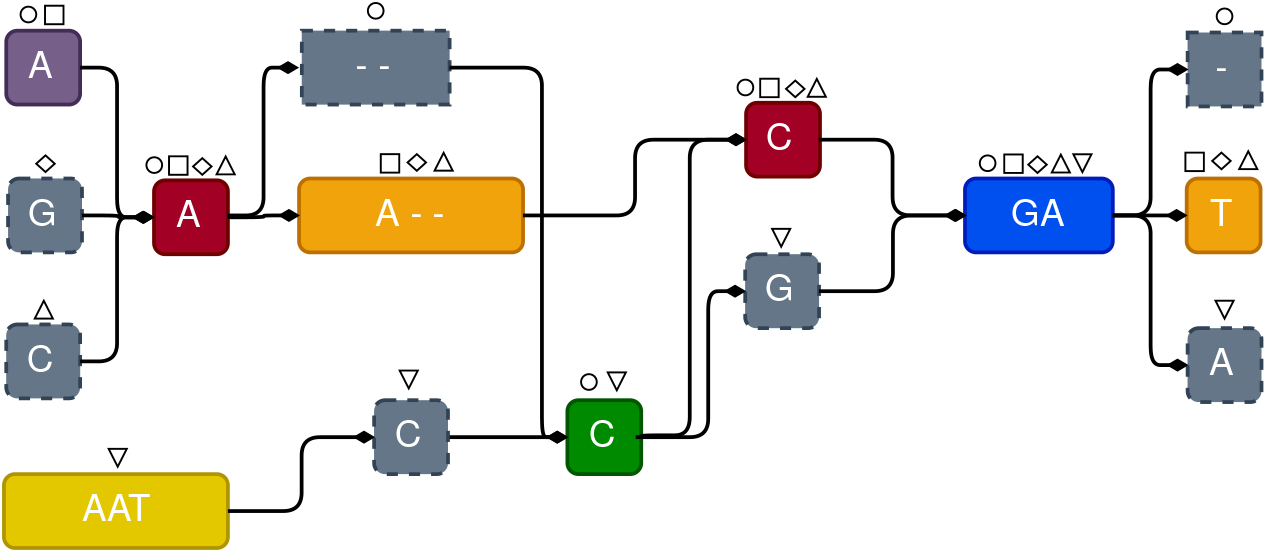
The Variation graph obtained from the Exact Cover of Figure 4, by defining a node for each block, and including arcs between consecutive blocks sharing at least one row in the MSA.

**Fig. 6:**
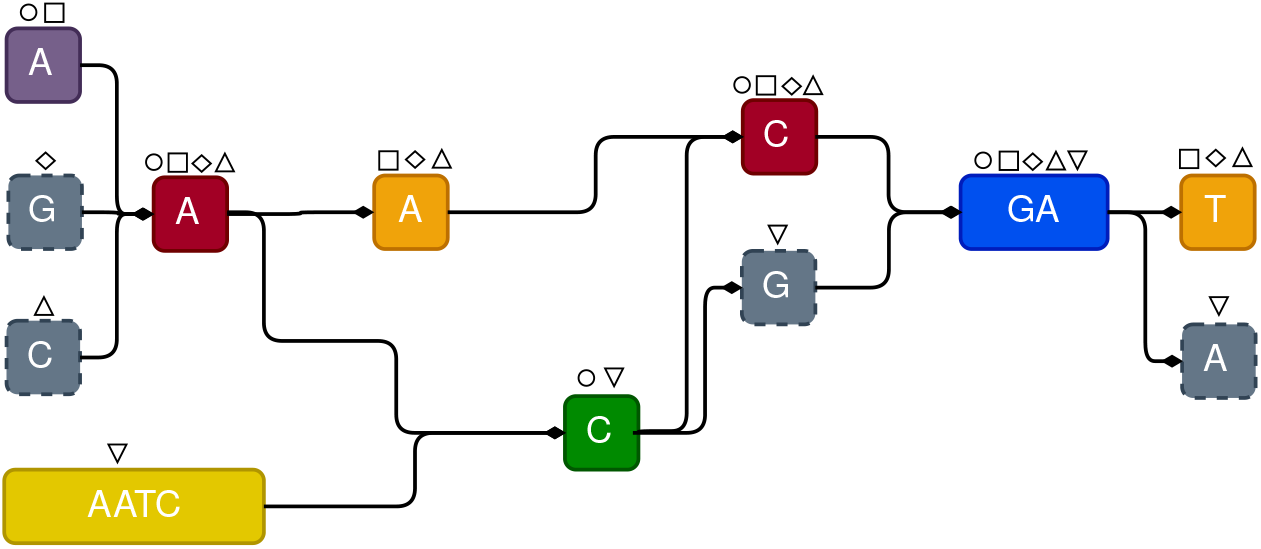
The result of postprocessing on the variation graph of Figure 5: indels are removed from node labels, and nodes labeled only by indels are removed. Non-branching paths are collapsed to a single node.

Finally, we collapse each non-branching subpath into a single node. Notice that the initial graph construction requires time linear in the MSA size, while the final step consists of a graph traversal that updates the graph. The creation of nodes and arcs from consecutive blocks is illustrated in Figures 4 and 5.

## 4 Solving the MWBC problem

Since the number of blocks of an MSA 𝒯 can be exponential in the MSA size, it is unfeasible to use a direct ILP formulation of the MWBC problem to obtain an optimal solution. Therefore we must describe some approaches that reduce the running time, based on restricting the set of possible blocks, that is solving an instance of MWBC-B, and forcing some blocks to be part of the solution.

We first look at blocks spanning all rows, called *vertical* blocks (see Figure 11). We introduce an additional parameter *α*: all maximal vertical blocks spanning at least *α* columns will be a block in our Exact Cover. The final effect is that each long substring that is shared by all rows of the MSA results in a single vertex of the variation graph, and all paths traverse such vertex. This heuristic will greatly reduce the running time, since we can split the MSA into smaller subMSAs, one for each portion delimited by two vertical blocks. For example, if a vertical block consists of columns 10–20 of the MSA and another vertical block consists of columns 57-81, and then we extract and solve the subMSA corresponding to the columns 21-56. Those subMSAs are then solved independently (if possible, in parallel) to obtain an exact cover of the initial ILP.

Even restricting to subMSA might result in a huge number of blocks, therefore we need to compute a set of blocks that is sufficiently small to be manageable but allows us to find an almost-optimal exact cover. To obtain such a set, we start from the ℬ set of all maximal blocks. Then a procedure, we called *decomposition*, is applied to each pair of overlapping maximal blocks to create some smaller blocks that are added to ℬ — we will detail in Section 4.2 the decomposition procedures.

Finally, we add some blocks to guarantee the existence of an exact cover in the set of blocks B, we will detail this procedure in Section 4.3.

### 4.1 Computing maximal blocks

As observed before, to obtain the instance ℬ for the MWBC problem, we first compute the set of *maximal* blocks, since their number is linear in the MSA size and they can be computed in linear time [22] via the Positional Burrows-Wheeler Transform (PBWT). In particular, in [22] the notion of maximal perfect haplotype blocks has been introduced in the framework of genome-wide selection. Maximal perfect haplotype blocks correspond to the notion of maximal block over a binary alphabet with the requirement that at the block has at least two rows. In our context, we used an implementation that extends the maximal haplotype block notion to the DNA alphabet extended with the indel symbol.

Still, an exact cover consisting only of maximal blocks might not exist, while we need to apply the ILP approach on a set ℬ of blocks that contains a feasible solution of the MWBC problem.

### 4.2 Decomposition of maximal blocks into non-overlapping blocks

First we will show (Lemma 1) that two overlapping maximal blocks can involve two nested intervals of columns if and only if the two sets of rows involved are one included in the other.

#### Lemma 1.

*Let* (*K*_1_, *b*_1_, *e*_1_) *and* (*K*_2_, *b*_2_, *e*_2_) *be two overlapping maximal blocks. Then b*_2_ ≤ *b*_1_ ≤ *e*_1_ ≤ *e*_2_ *if and only if K*_2_ ⊂ *K*_1_.

*Proof*. (⇒) Assume to the contrary that ∃*r* ∈*K*_2_\ *K*_1_. Since *b*_2_ ≤ *b*_1_ ≤ *e*_1_ ≤ *e*_2_, this implies that *r*[*b*_1_ : *e*_1_] is equal to the label of the block (*K*_1_, *b*_1_, *e*_1_), hence contradicting the maximality of (*K*_1_, *b*_1_, *e*_1_).

(⇐) Assume now that *K*_2_⊂ *K*_1_. Since the two maximal blocks are overlapping, *b*_2_ ≤ *e*_1_ ≤*e*_2_ or *b*_2_ ≤ *b*_1_ ≤*e*_2_. Assume that *b*_2_ ≤*e*_1_ ≤ *e*_2_ (the other case is symmetrical), and assume to the contrary that *b*_1_ *< b*_2_. Since *K*_2_⊂ *K*_1_, also (*K*_2_, *b*_1_, *e*_2_) is a block, contradicting the maximality of (*K*_2_, *b*_2_, *e*_2_).

The coverage of an MSA can be split into two parts: those positions covered by maximal blocks and the remaining ones. In the latter case, there exists a unique option to cover the positions, that consists in creating one-row blocks. We can now focus our attention on finding the coverage of the regions covered by maximal blocks.

Our purpose is to define a procedure to create blocks contained in maximal blocks. We introduce the first notion of *decomposition* called row-maximal decomposition, which builds on Lemma 1. It creates blocks from two intersecting maximal blocks such that each new block preserves all rows (its row-maximality) of the maximal block it is contained in.

#### Definition 8

(Row-maximal decomposition of maximal blocks). *Given two overlapping blocks l*_1_ = (*K*_1_, *b*_1_, *e*_1_) *and l*_2_ = (*K*_2_, *b*_2_, *e*_2_) *with b*_1_ ≤ *b*_2_, *the result of their row-maximal decomposition consists of the following blocks:*

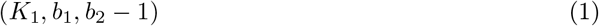

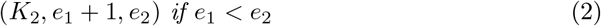

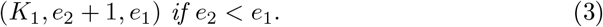

Notice that more than one condition might be true in Definition 8. In that case, the result consists of all blocks whose conditions hold. We show two examples of rowmaximal decomposition in Figure 7, where we can observe that all columns part of the intersection are excluded.

**Fig. 7:**
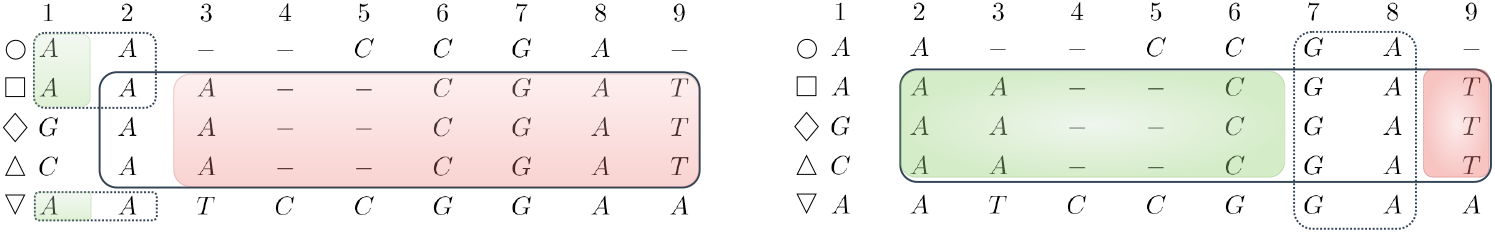
The additional blocks obtained by the *row-maximal decomposition* on the overlapping maximal blocks of Figure 2 (on the left) and of Figure 3 (on the right).

The case when *b*_2_ *<* 1 is symmetrical to the one of Definition 8.

A second decomposition, called complete decomposition, extends the row-maximal decomposition, by creating blocks on the shared columns of the intersection of maximal blocks, these new blocks are fully contained in only one maximal block or in both at the same time, see an example in Figure 8.

**Fig. 8:**
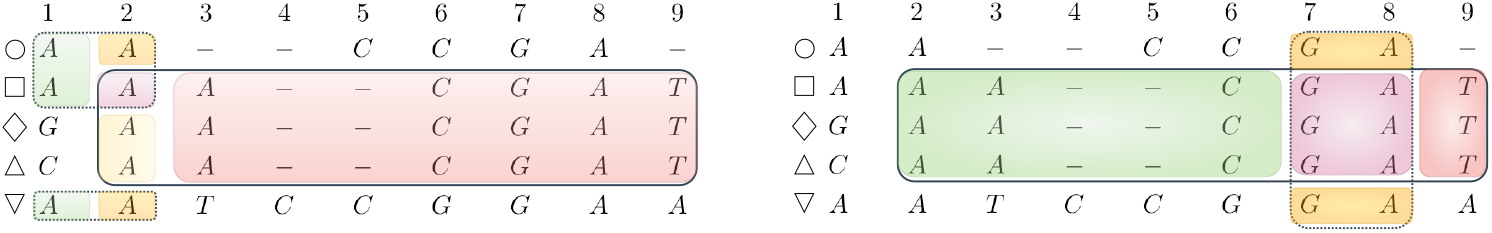
The additional blocks obtained by the *complete decomposition* on the overlapping maximal blocks of Figure 2 (on the left) and of Figure 3 (on the right). Additional blocks are colored.

This second decomposition returns a superset of the blocks computed with a row-maximal decomposition, hence resulting in a larger set of blocks. Just as for Definition 8, we describe only the case when *b*_1_ ≤ *b*_2_, since the other case is symmetrical.

#### Definition 9

(A complete decomposition of maximal blocks). *Given two overlapping blocks l*_1_ = (*K*_1_, *b*_1_, *e*_1_) *and l*_2_ = (*K*_2_, *b*_2_, *e*_2_) *with b*_1_ ≤ *b*_2_, *the result of their complete decomposition consists of the union of the result of their row-maximal decomposition, and of the following blocks:*

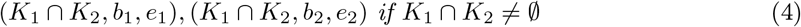

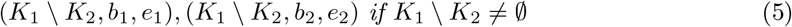

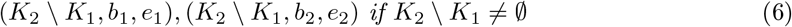

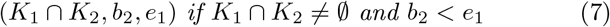

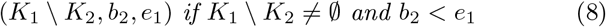

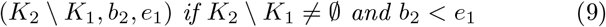

*Notice that more than one of the above conditions might be true. In that case, the result consists of all blocks whose conditions hold*.

For each pair of overlapping maximal blocks, we compute their decomposition and the result is added to the set of blocks. In Section 5 we will discuss when to use the row-maximal or complete decomposition.

### 4.3 Adding short blocks

Maximal blocks involving only one row — blocks (*K, b, e*) with |*K*| = 1 — correspond to vertices of the variation graph encoding an entire genome, without highlighting the common portions of the genomes we want to represent in a pangenome. For this reason, they are undesirable and we remove them from the set ℬof blocks. On the other hand, their removal does not guarantee that ℬ contains an exact cover. To overcome this problem, we add to ℬ all blocks of the form ({*r*}, *b, e*) where [*b, e*] is a maximal interval in the row *r* not covered by any maximal block spanning at least two rows. We refer to these blocks as *one-row blocks*.

Finally, we add to ℬ all blocks of the form (*K*_*σ*_, *b, b*) for each 1 ≤ *b* ≤ *n* and for each character *σ*, where *K*_*σ*_ is the set of rows that have the character *σ* in column *b*. Notice that, for each column *b* there are at most |Σ| + 1 such blocks, since the character *σ* can be the indel. We refer to these blocks as *one-character blocks*. The addition of one-row and one-character blocks guarantees that the set ℬcontains a feasible solution of MWBC.

### 4.4 An ILP for MWBC

We can now describe the ILP for solving the MWBC that forms the main ingredient of our approach. We recall that the set of blocks ℬ fed to the solver consists of the following blocks: i) the set of maximal blocks, ii) the result of the decomposition of the maximal blocks described in Section 4.2 (which is either the row-maximal or the complete decomposition), iii) the one-row blocks, and finally iv) the one-character blocks described in Section 4.3. Our implementation also removes duplicate blocks, but that is not relevant to our discussion.

The binary variables we use are *C*[*K, b, e*] for each block (*K, b, e*), where *C*[*K, b, e*] = 1 iff the block (*K, b, e*) belongs to the solution. The main constraints in our ILP are the usual ones used to encode an exact cover:

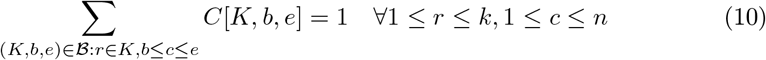

The above constraints express the condition that exactly one block in the solution covers each position (*r, c*) of the MSA.

We have not introduced the objective function of our ILP approach, since different biological or computational goals can be better modeled with different objective functions: one of the advantages of an ILP approach is that we do not rely on a specific objective function, but we can leave it as a decision that the final user will take.

We recall that each block in the solution will be later transformed into a node of a Variation Graph. We are going to show five possible objective functions, the first three will be a subject of our experimental assessment in Section 5.

We will denote by *γ*(*K, b, e*) the label of the blocks (*K, b, e*) without considering indels. Clearly, the length of *γ*(*K, b, e*), denoted as |*γ*(*K, b, e*)|, is at most *e* − *b* + 1 and it is equal to *e* − *b* + 1 when the string in the block does not contain indels.

1. Minimize the number of blocks. The objective function is called **blocks** and it is defined as

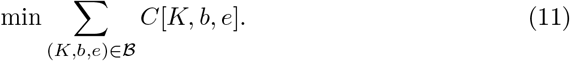
2. Minimize the number of vertices of the variation graph, penalizing vertices whose labels are shorter than a certain threshold *q*. Notice that read mappers based on seeding, such as GraphAligner [14], can benefit from graphs with longer node labels, where it is more likely to find a seed. The objective function is called **weighted** and it is defined as

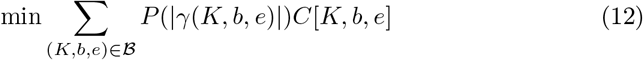

where the penalization function *P* is defined as

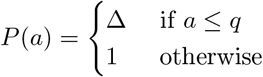

and Δ *>* 1 is the penalization parameter for these shorter labels.
3. Minimize the number of vertices of the variation graphs, penalizing vertices that are not used by at least *p* input sequences, that is we prefer nodes with high coverage. The rationale is that some variant calling pipelines (such as minigraph-cactus or `vg` prune) start by removing vertices with low coverage. The objective function is called **depth** and it is defined as

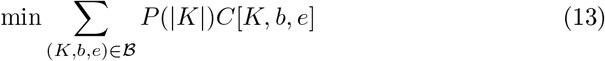

where the penalization function *P* is

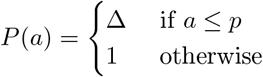

with Δ *>* 1 the penalization parameter.
4. Minimize the total length of the labels of the nodes of the variation graph, excluding indels. The objective function is called **strings** and it is defined as

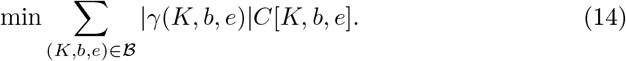
5. Minimize the total length of the labels of the variation graphs, penalizing vertices that are shared by fewer sequences. The objective function is called **penalized strings** and it is defined as

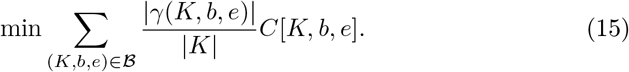

## 5 Experimental analysis

We randomly select 50 complete SARS-CoV-2 genome sequences from ENA and downloaded them with `ENA Tools`^2^, we then create an MSA using MAFFT (version 7.525) with default parameters. For results with 20 and 100 sequences see [Additional file 1]. Our analysis is divided into 3 parts. The first part examines the scalability of the decomposition strategy, by determining the time and memory needed to apply the *row-maximal* and the *complete* decomposition on the input MSAs. Secondly, we analyze the effect of varying the parameter *α*, that is the minimum length of forced vertical blocks. More precisely, we report the values of the objective function *blocks* for different values of *α*, using both decompositions. Finally, we study the effect of choosing different objective functions on the resulting variation graph. The graphs are evaluated on the number of vertices, the length of the graph labels, the number of potential seeds it contains, and the number of shared nodes they have. In this last part, we have compared the variation graphs computed by `pangeblocks` with those created with `vg`, `make_prg`, `founderblockgraph` (starting from the same MSA), and `pggb` (using the same set of sequences).

While our approach requires more computational resources, the experiments show that `pangeblocks` can be run on a personal computer for real instances.

### 5.1 Assessing the decomposition strategies

We start by investigating the impact of the decomposition strategies on the computational resources required to run `pangeblocks`. Since the blocks obtained with the complete decomposition (Definition 9) include those obtained with the row-maximal decomposition (Definition 8), selecting the complete decomposition leads to a larger ILP instance.

We ran `pangeblocks` varying the number of columns, ranging from 100 to 3000 (incrementing by 100). For each number of columns, we have extracted 10 random MSAs from the complete MSA on 50 SARS-CoV-2 genomes. We ran all those experiments on the *blocks* objective function, since its the choice should not affect the memory and time, and without any forced vertical block (this is imposed by setting *α* equal to the length of the MSA).

RAM usage and user time are comparable when using both decompositions in instances up to a few hundred columns. After that point, the complete decomposition requires more time and memory than the row-maximal decomposition (see Figure 9). In fact, the complete decomposition failed since it used too much RAM (more than 90GB) on approximately 18% of the instances — only successful runs are displayed in the Figure.

**Fig. 9:**
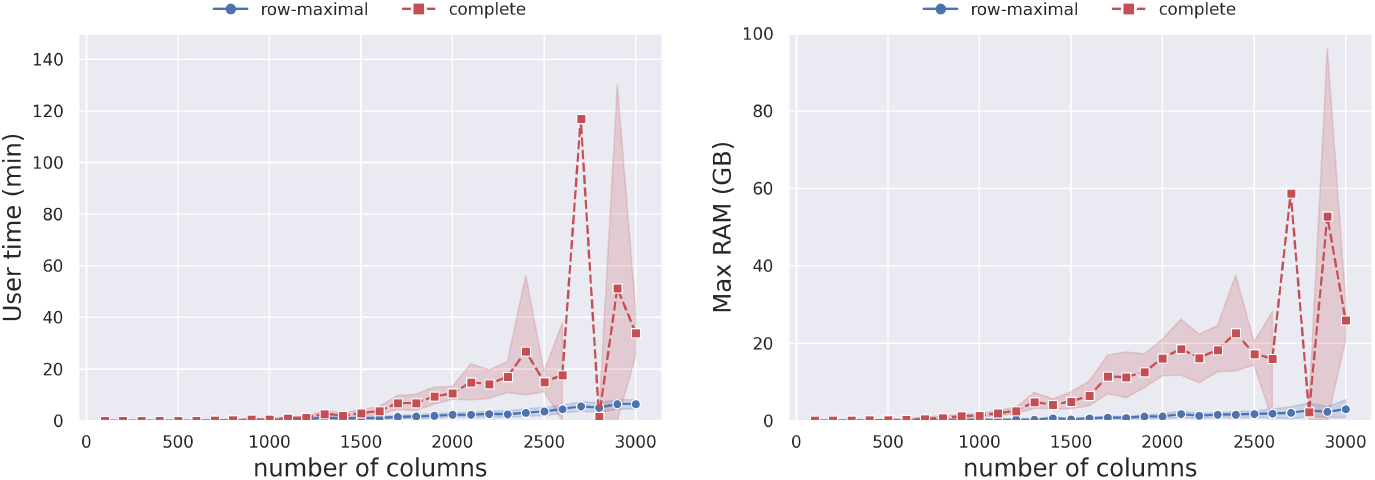
Time (on the left) and peak RAM usage (on the right) needed to solve the ILP, as a function of the MSA length and the choice of decomposition. For each MSA length, ranging from 100 to 3000, incremented by 100, we have extracted 10 random MSAs with that length from the MSA of 50 SARS-CoV-2 complete genomes. The orange dashed line represents the complete decomposition, and the blue line represents the row-maximal decomposition. The objective function *blocks* was used in this experiment.

Notice that the row-maximal decomposition never exceeded 12 GB of RAM and completed its execution in less than 10 minutes.

### 5.2 Assessing the choice of *α*

The experiment in this section is run on the MSA on 50 complete SARS-CoV-2 genomes. This MSA has 29903 columns.

We recall that the parameter *α* is the minimum number of columns of all forced vertical blocks, that is all vertical blocks spanning at least *α* columns are forced to be part of the solution. We explore if vertical blocks are chosen by `pangeblocks` with different objective functions.

We focus on those *α* values where the longest subMSA (between two forced vertical blocks) has more columns than the ones with smaller values of *α*, we refer to those values of *α* as *breakpoints*. Breakpoints are especially relevant since each subMSA can be solved independently and in parallel, therefore the computational resources needed are affected mostly by the size of the largest subMSA (*i*.*e*. the one that changes at a breakpoint). Breakpoints are shown in Figure 10 and can be identified as the values of *α* where the blue curve increases its value. The effect of choosing different values of *α* on the entire MSA can be seen in Figure 11, where black rectangles represent the forced vertical blocks, white areas are the subMSA with the largest number of columns, and the gray rectangles are all other subMSAs.

**Fig. 10:**
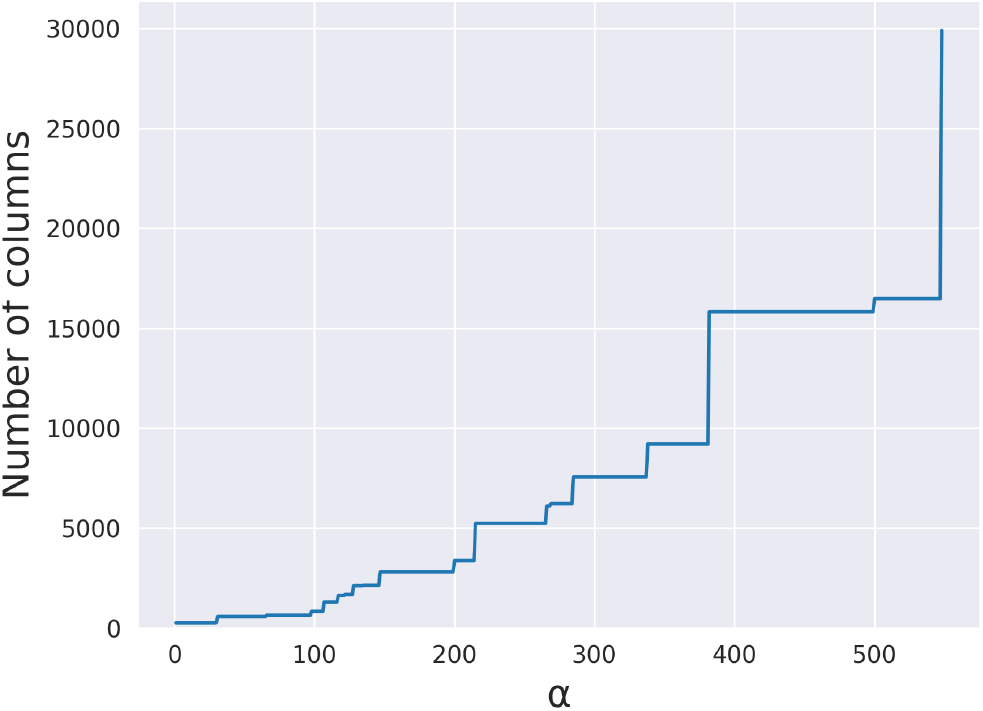
Longest (columns) subMSA obtained for different values of *α* in the SARSCoV-2 MSA with 50 sequences and 29903 columns. The number of columns (y-axis) shows the longest subMSA given that all vertical blocks with length at least *α* (xaxis) are fixed. The curve shows how increasing the value of *α* implies an increase of the number of columns in the longest subMSA. In particular, we call *breakpoints* of *α* those values of *α* that determines a change in length of the subMSA. This plot can help to decide a reasonable value of *α* given the available resources. A picture of the MSA given these breakpoints is shown in Figure 11.

**Fig. 11:**
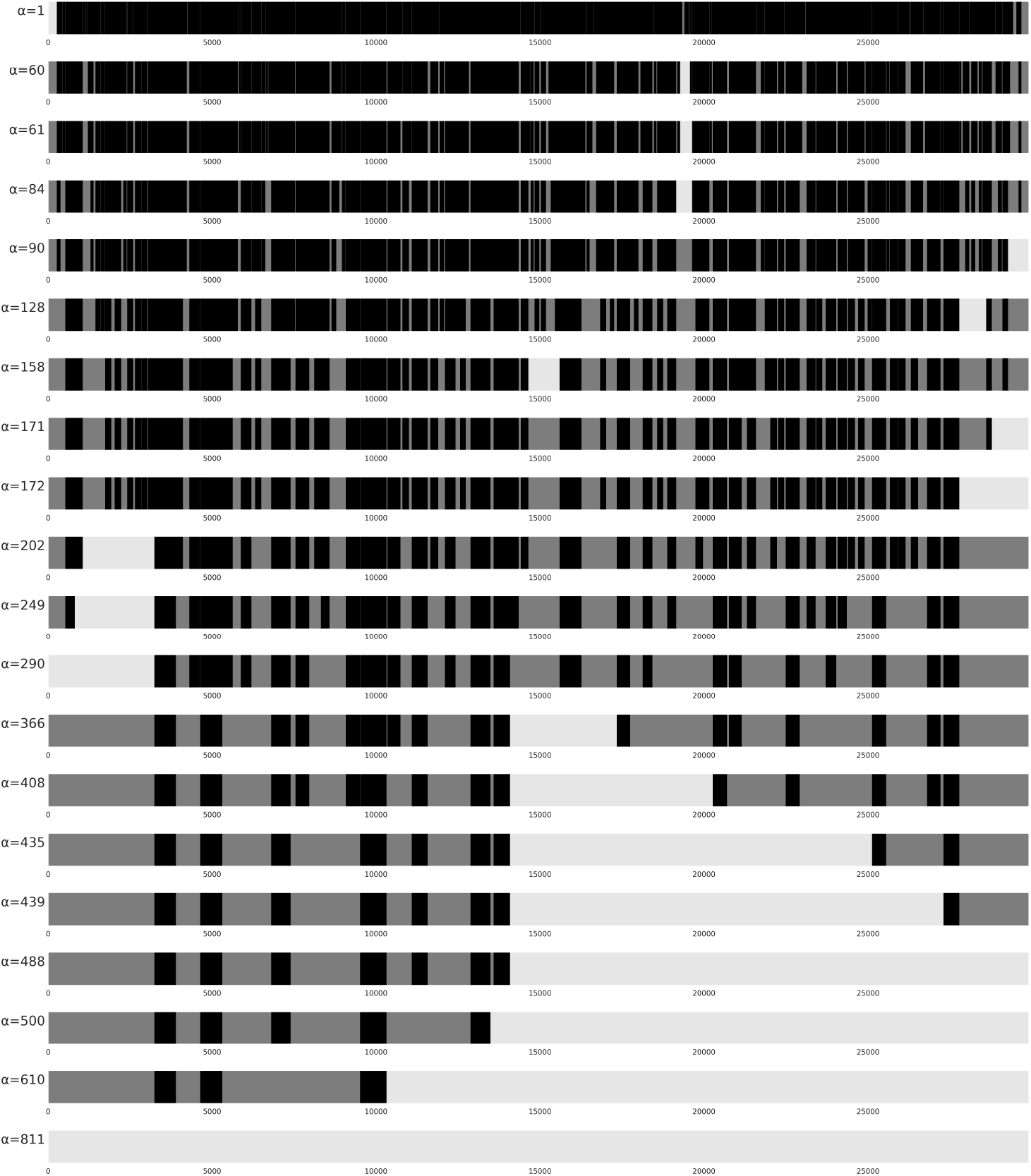
A view of SARS-CoV-2 MSA with 50 sequences and 29903 columns for different values of *α* (see label to the left), from *α* = 1 to *α* = 881 (from top to bottom). Forced vertical blocks (those with the number of columns less or equal to *α*) are shown in **black**. SubMSAs in between vertical blocks (or at the beginning, or the end of the MSA) are in <u>gray</u>. The longest subMSA is shown in <u>light gray</u>. Values of *α* correspond to *breakpoints* shown in Figure 10, where the longest subMSA when fixing *α* increases in length.

Notice that we have decided to focus on the number of columns of the largest subMSA instead of the time and memory used, since Section 5.1 shows that the complete decomposition can manage at most a few thousand columns using at most 90 GB of RAM. On the other hand, Figure 10 represents all possible number of columns of a subMSA, hence it does not have such restriction. Moreover, each subMSA is solved independently, therefore we can actually use the number of columns of the largest subMSA as a proxy of the computational resources needed.

Now we can analyze the effect of the choice of *α* on the number of vertices of the resulting graph — results are shown in Figure 12 — when we use the *blocks* objective function. Since we need to run `pangeblocks` on the resulting subMSAs, we have been able to consider only values of *α* smaller than 158.

**Fig. 12:**
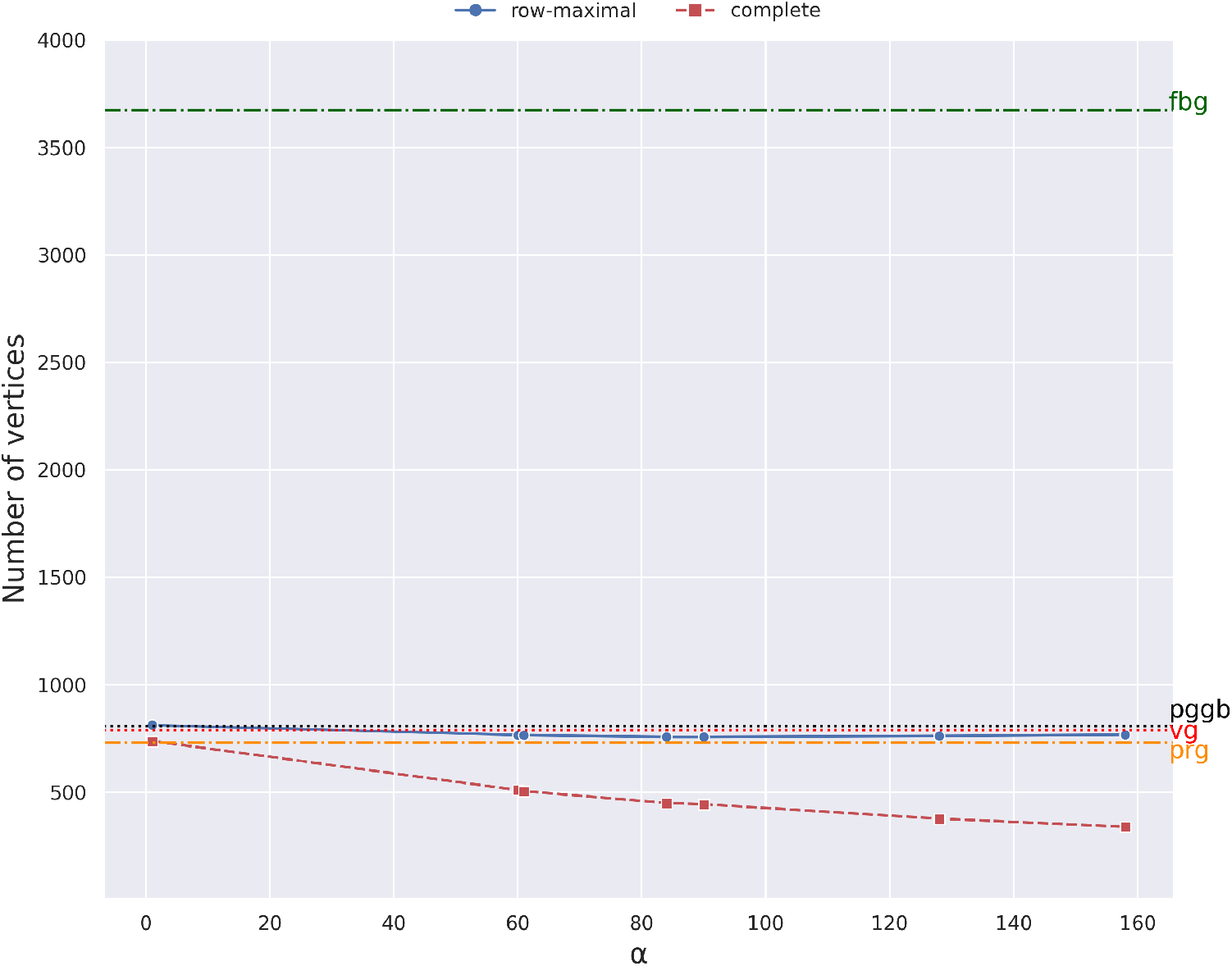
Number of vertices. The plot is the results of running `pangeblocks`, `pggb`, `vg`, `make_prg` and `founderblockgraph` on the 50 SARS-CoV-2 genomes instance, using the row-maximal (blue dot) and the complete (red square) decomposition. The y-axis is the number of vertices of the variation graphs (after post-processing), and the x-axis corresponds to various values of *α* (from 1 to 158, see Figure 11). The number of nodes for other tools is shown by horizontal lines. The black dotted line corresponds to `pggb`, the red dashed line corresponds to `vg`, and the orange dash-dot line corresponds to `make_prg`, and the green dash-dot line corresponds to `founderblockgraph`.

We expected larger values of *α* to correspond to fewer vertices, but this is what we observe for the complete decomposition only, and even in that case it is not a monotonous decrease. This phenomenon is due to the fact the maximal blocks and their decomposition is computed independently for each subMSA, therefore the set of blocks fed to the ILP for larger values *α* is not a superset of the set for smaller values.

Moreover, the phase where a solution of MWBC is used to compute a variation graph introduces an additional uncertainty in the number of vertices of the final graph.

We observe in Figure 12 that the row-maximal decomposition reach a plateau, or even begin to worsen, for moderate values of *α* — around 90 for the row-maximal decomposition — hinting that the large computational resources needed for larger values of *α* are not actually necessary in this case. In the case of the complete decomposition, due to computational resources, we are limited to consider up to *alpha* = 158, nevertheless, in smaller instances (See Supplementary material) we observe the same behavior described for the row-maximal decomposition.

It is especially noticeable that both decompositions are able to obtain graphs with fewer vertices than `vg` and `pggb` as shown in Figure 12. This is partially surprising since `vg` and `pggb` can build variation graphs that contain cycles (which helps in reducing the size of the graph), while `pangeblocks` constructs only acyclic graphs. `make_prg` creates a graph with the number of nodes in between both decompositions, while `founderblockgraph` graph has more than 3× the number of nodes than all other tools. The row-maximal decomposition results in essentially the same number of vertices as `vg` and about 10% fewer vertices than `pggb`. On the other hand, the more computationally expensive complete decomposition obtains 55-60% fewer vertices than `vg`.

By choosing *α* we are fixing vertical blocks with equal or more columns than *α*, this imposes finding a suboptimal solution of the MSA by solving an ILP for each resulting subMSA not covered by the fixed vertical blocks. Vertical blocks are nodes in the graph traversed by all sequences. We can see in Figure 13 (right) that with the complete decomposition, the *blocks, weighted*, and *depth* objective functions lead to fewer nodes that are shared by all sequences when *α* is increased. With larger values of *α*, the ILP can discard (unforced) vertical blocks to find a better solution that contains larger blocks, i.e. those that cover more cells in the MSA. In other words, both the number of blocks and the total length of labels of the graph decrease with larger values of *α*, as expected since the space of the feasible solution is larger.

**Fig. 13:**
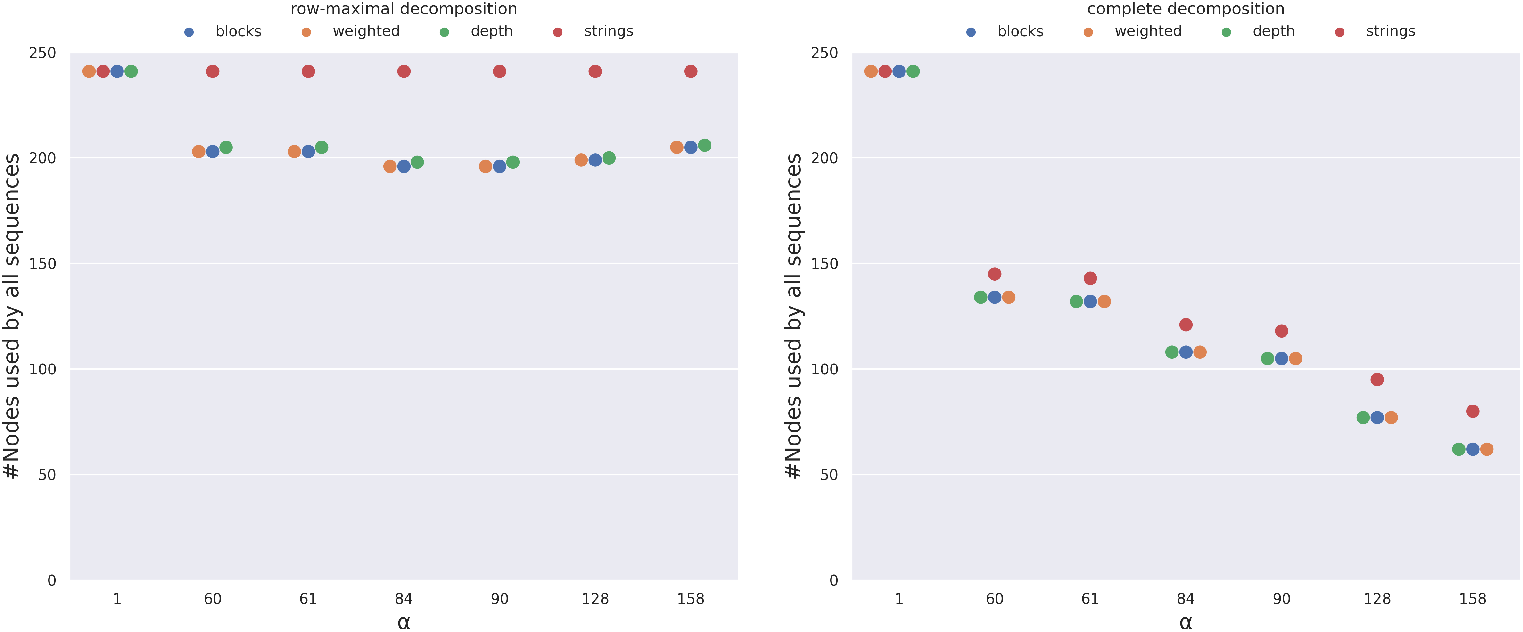
Number of vertices of the variation graph that are traversed by all sequences as a function of *α* and the choice of decomposition, on the MSA of 50 SARS-CoV2 complete genomes. For the complete decomposition (right), the objective functions *blocks, weighted*, and *depth* coincide in the use of vertical blocks (y-axis), and decrease when increasing *α*, the same tendency for *strings* with higher usage of vertical blocks.In the row-maximal decomposition (left), the tendency is to decrease the use but with lower fluctuation due to the smaller search space. The *strings* objective function does not change with *α*, meaning that vertical blocks are always preferred.

The left plot of Figure 13 is on row-maximal decomposition, where there is no substantial difference among the objective functions. When choosing the *depth* objective function, the procedure prefers to pick blocks traversed by a larger number of sequences. Among those, all vertical blocks fit this criteria, therefore they are likely to be retained even for larger values of *α*, hence the final solution computed is largely unaffected by the choice of *α*.

This is likely to be the result of the smaller search space created by this decomposition, which is heavily constrained and does not allow the choice of the objective function to affect the selection of the blocks covering the MSA. On the other hand, larger values of *α* and the associated larger search space lead to fewer nodes overall, including those shared by all nodes. Finally, comparing the two plots of Figure 13 we can observe that the two decompositions lead to a completely different behavior of the ILP when choosing the different objective functions, except for *α* = 1, when all vertical blocks are forced to be in the solution.

### 5.3 Assessing the objective functions

In this section, we are going to assess the effect of the objective functions on some properties of the resulting variation graph. More precisely, we are going to analyze the blocks, weighted, and depth objective functions, since the strings and the penalized strings objective functions result in graphs that have a simplistic structure. The optimal solution with the strings objective function consists of one-character blocks, therefore demanding the nontrivial graph optimization steps to the post-processing phase.

Figure 14 (a) shows the total number of vertices of the variation graphs (after postprocessing) obtained with the different objective functions, with *α* = 168. In all three settings, the number of vertices is smaller than those achieved by `pggb` and `vg`, even though the difference with `vg` is negligible when using the row-maximal decomposition. Choosing the *blocks* or the *weighted* objective functions produces graphs with the fewest vertices.

**Fig. 14:**
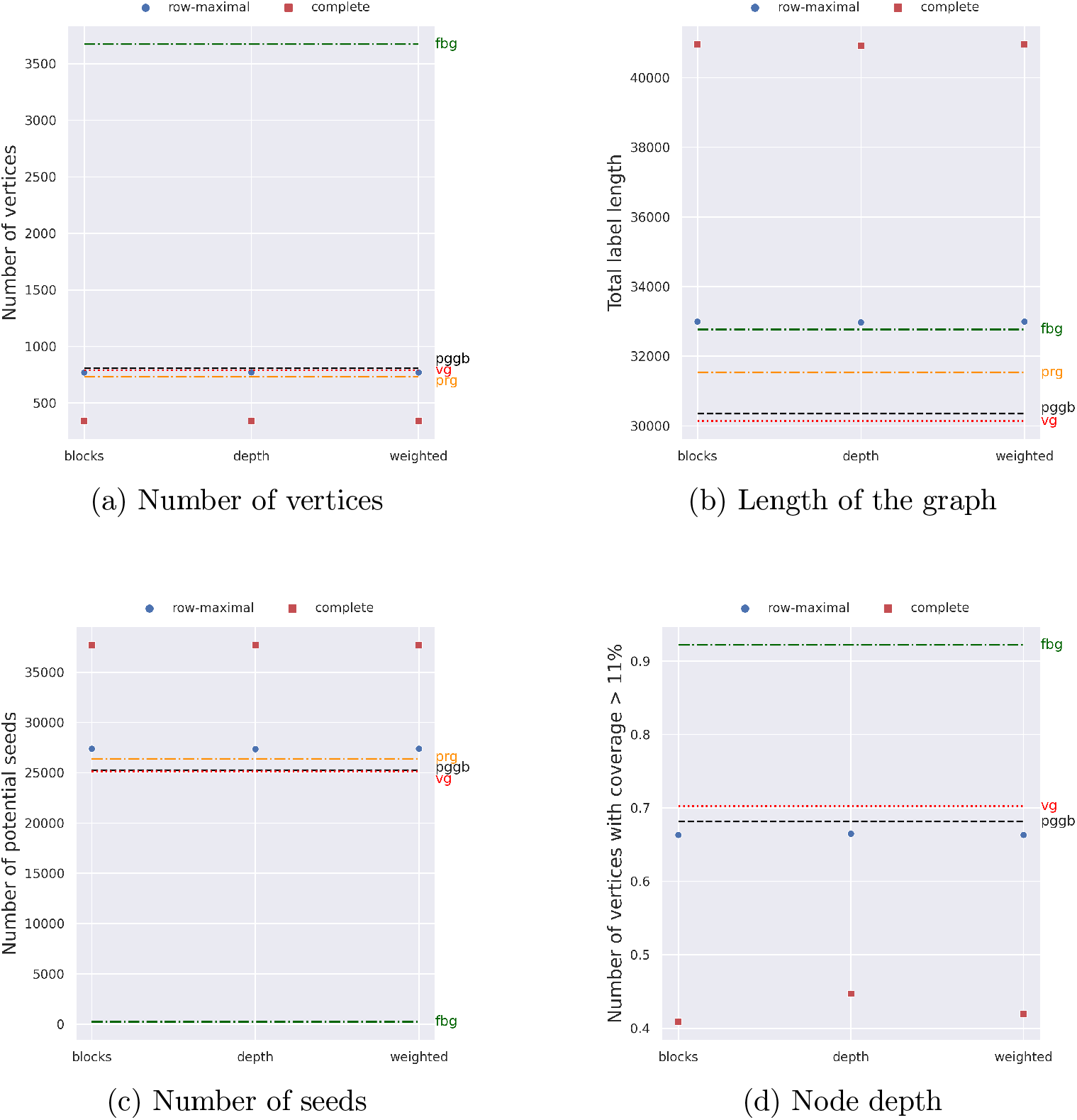
Four metrics were measured over pan-genome graphs created with five construction tools on the 50 SARS-CoV-2 genome instance. We report (a) the number of vertices of each graph, (b) the total length of the graph, (c) the number of potential seeds (k-mers) of length 20, and (d) the number of vertices used by at least the 11% of the input sequences. For `pangeblocks` we report the results of three objective functions: *blocks, depth*, and *weighted* (horizontal axis) The metrics for other tools are shown by horizontal lines. The black dotted line corresponds to `pggb`, the red dashed line corresponds to `vg`, the orange dash-dot line corresponds to `make_prg`, and the green dash-dot line corresponds to `founderblockgraph`. Since `make_prg` creates a sequence graph, node depth cannot be computed for it.

When choosing the weighted objective function, we have run `pangeblocks` with the parameter *q* equal to 20 (all blocks spanning fewer than *q* columns have cost Δ = 1000 instead of 1). We present only the results with Δ = 1000, since smaller values of Δ result in graphs that are very close to those obtained with Δ = 1, that is without penalization. When choosing the depth objective function, we have run `pangeblocks` with the parameter *p* equal to 0.11 (all blocks that are traversed by fewer than a *p* fraction of the input sequences have cost Δ = 1000 instead of 1). When choosing the blocks objective function, no parameter is needed.

Notice that choosing the *weighted* objective function results in a graph with negligible differences from the one created by choosing the *blocks* objective function.

We recall that the objective function determines the block cover, while the *y* axis of the figure is computed on the result of the post-processing over the block cover, nodes that are labeled only by indels are removed. The *weighted* objective function is designed to prefer nodes with longer labels, by penalizing those who contain too many indels, see Eq. 12, which leads to the selection of fewer, but longer (w.r.t. their label without indels) nodes, as we can see in Figure 14 (b).

Figure 14 (b) shows the total number of characters of the graph labels (after postprocessing) obtained with the different objective functions, with *α* = 158. In all three settings, the total number of characters obtained by `pangeblocks` is larger than that achieved by `pggb` and `vg`, even though the differences are marginal when using the row-maximal decomposition.

Figure 14 (c) shows the total number of potential seeds (that is the 20-mers that are substring of at least a vertex label) of the variation graphs, obtained with the different objective functions, with *α* = 158.

When using the complete decomposition, we obtain a number of seeds that is larger than those achieved by `pggb` and `vg`. When choosing the row-maximal decomposition, the results obtained by `pangeblocks` are marginally larger than both `vg` and `pggb`. The *weighted* objective function produces a graph with more seeds, thus the graph can be potentially more informative if the task is aligning reads to the graph.

Figure 14 (d) shows the fraction of vertices traversed by at least 11% of the genome sequences, obtained with the different objective functions, with *α* = 158. When using the row-maximal decomposition, the results are slightly smaller than `pggb`. `vg` on the other hand has the highest number of these nodes. In the case of `pangeblocks`, we notice that choosing the *depth* objective function results in a higher number of vertices with this coverage, for both decompositions, as was expected, since this objective function was built for this purpose.

In general, we can observe that `founderblockgraph` creates bigger graphs w.r.t. the other tools, both in number of nodes and length of the graph, with the particularity of very short nodes, which translates into a small number of potential seeds. `make_prg` creates graphs with fewer nodes than `vg`, `pggb`, and `founderblockgraph`, but with longer nodes, showing a higher number of potential seeds than the other tools.

## 6 Conclusions and future directions

We have presented a computational problem, Minimum Weight Block Cover (MWBC), that formalizes the problem of constructing a variation graph from an MSA. We have proposed an ILP approach for solving the MWBC problem and constructing a graph and we implemented it as a tool, called `pangeblocks`.

We have experimentally shown that `pangeblocks` can scale to genome-scale instances. More precisely, our experimental analysis has been run on an MSA on 50 complete SARS-CoV-2 genome sequences (approximately 30kbp). We have also shown how the ILP formulation is sufficiently flexible to accommodate different goals. In fact, by choosing an objective function, the decomposition strategy, or the minimum size of a forced vertical block, we are able to obtain variation graphs that have different characteristics, for example we can obtain graphs whose number of vertices ranges from 800 to 1400. Notice that all graphs computed by `pangeblocks` are complete representations of the input MSA, that is each input genome corresponds to the concatenation of the labels if a path of the graph.

We have compared `pangeblocks` with the two most widely used tools for constructing variation graphs, `vg` and `pggb`, showing that we can obtain graphs that are similar to theirs. We have to remind that our ILP approach is more flexible, therefore allowing to tailor the graph construction to specific need. At the same time, those possibilities affect the computational resources needed by `pangeblocks`.

We observe that the use of ILP and optimization criteria in the construction of the variation graphs allows to include biological constraints. In particular, we have introduced the parameter *α* to bound the length of the so called vertical blocks, i.e. those nodes of the graph corresponding to consecutive MSA columns where the input sequences are all identical. While we have introduced *α* for computational reasons, setting its value might have a possible biological motivation, since it can influence the regions where recombinations are more likely.

There are some directions for future research. First of all, there is a large gap in both the computational resources needed and the resulting graphs, between the complete and the Row-maximal decompositions. Therefore we think that different decomposition strategies can be relevant. Moreover, we would like to keep most of the flexibility allowed by the ILP approach, but avoiding its computational cost. This requires restricting the possible parameters to a subset of those that are currently modeled by the ILP, and designing a combinatorial approach that is able to quickly compute a solution on those parameter.

## 7 Declarations

## 7.1 Abbreviations

ENA: European Nucleotide Archive.
G-MWBC: General Minimum Weight Block Cover.
ILP: Integer Linear Programming.
MSA: Multiple Sequence Alignment.
MWBC: Minimum Weight Block Cover.
NGS: Next-Generation Sequencing.
PBWT: Positional Burrows-Wheeler Transform.

## 7.2 Ethics approval and consent to participate

Not applicable.

## 7.3 Consent for publication

Not applicable.

## 7.4 Availability of data and materials

- Project name: `pangeblocks`
- Project home page: https://github.com/AlgoLab/pangeblocks [26]
- Archived version: doi.org/10.5281/zenodo.12187814
- Operating system(s): Platform independent
- Programming language: C++, Python
- Other requirements: gurobi 9.5.2, snakemake>=7.22.0
- License: GPL-3.0

## 7.7 Author’s contributions

JAC, PB, GDV: Conceptualization, Methodology, Writing - Original Draft. JAC, GDV: Software. JAC, GDV, LD, SC: Data Curation, Validation. JAC, PB, GDV, LD, SC: Writing - Review and Editing.

## 7.9 Acknowledgments

Not applicable.

## Additional Material

- File name: Additional file 1
- File format: Additional file 1.pdf
- PangeBlocks: customized construction of pangenome graphs via maximal blocks. Supplementary Material.
- Additional experiments with 20 and 100 sequences.

## Footnotes

1 github.com/AlgoLab/Wild-pBWT [23]

2 https://github.com/enasequence/enaBrowserTools

