## Additional file 1 for "PangeBlocks: customized construction of pangenome graphs via maximal blocks"

[1]DISCo, Univ. Milano – Bicocca, Viale Sarca, Milano, 20126, Italy.

Contributing authors:;

**Keywords:** pangenome graphs, variation graph construction, integer linear programming, maximal blocks

#### 1 SARS-CoV-2 MSA with 20 sequences

We report the same results shown in the main manuscript for the SARS-CoV-2 MSA, this time with 20 sequences.

A view of SARS-CoV-2 MSA with 20 sequences is shown in 3, where black colors represent vertical blocks of length at least $\alpha$ (left axis) fixed to the final solution. Gray (and white) areas correspond to portions of the MSA fed to the ILP. White areas correspond to the longest subMSA.

The longest (in number of columns) subMSAs depend on the $\alpha$ that is fixed, values of $\alpha$ where the longest subMSA increases its value are called breakpoints, and are shown in 4.

RAM usage and User time are shown in 1.

The number of vertices, length of the graph, number of potential seeds, and number of vertices with coverage $> 11\%$ are shown in 5, 6, 7, 8, respectively. These values are shown using $\alpha = 659$.

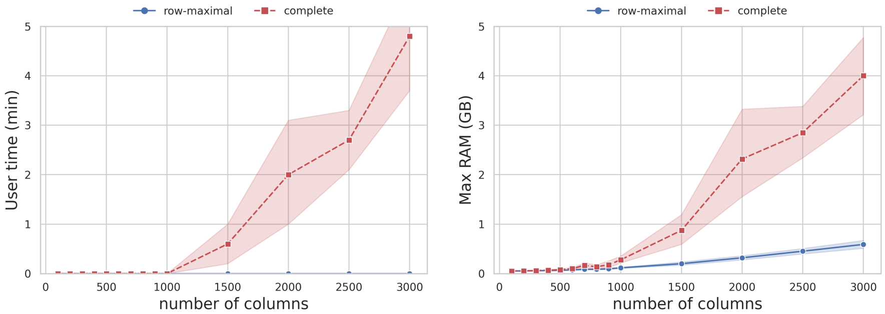


**Fig. 1**: Time (on the left) and peak RAM usage (on the right) needed to solve the ILP, as a function of the MSA length and the choice of decomposition. For each MSA length, ranging from 100 to 3000, incremented by 100, we have extracted 10 random MSAs with that length from the MSA of 20 SARS-CoV-2 complete genomes. The red dashed line represents the complete decomposition, and the blue line represents the row-maximal decomposition. The objective function *blocks* was used in this experiment.

### 2 SARS-CoV-2 MSA with 100 sequences

A view of SARS-CoV-2 MSA with 100 sequences is shown in 9, where black colors represent vertical blocks of length at least $\alpha$ (left axis) fixed to the final solution. Gray (and white) areas correspond to portions of the MSA fed to the ILP. White areas correspond to the longest subMSA.

The longest (in the number of columns) subMSAs depend on the $\alpha$ that is fixed, values of $\alpha$ where the longest subMSA increases its value are called breakpoints, and are shown in 10.

Results are only reported with $\alpha = 1$. In table 1 is reported the RAM usage and User time. These values correspond to the average and standard deviation of running `pangeblocks` with all five objective functions with the same parameters used in the main manuscript.

The number of vertices, length of the graph, number of potential seeds, and number of vertices with coverage $> 11\%$ are shown in 11, 12, 13, 14, respectively.

**Table 1**: `pangeblocks` **Maximum RAM usage and User Time on the 100 SARS-CoV-2 MSA** Results for 100 sequences are obtained using $\alpha = 1$ and all objective functions, both were run with the same parameters used in the main experiments. We report the average and standard deviation for each decomposition. Max RAM is in GB, and User Time is in seconds.

| | row-maximal | | complete | |
|---|---|---|---|---|
| Dataset | User Time | RAM | User Time | RAM |
| 100 | $63.45 \pm 3.85$ | $0.72 \pm 0.007$ | $889.65 \pm 59.10$ | $11.776 \pm 0.0008$ |

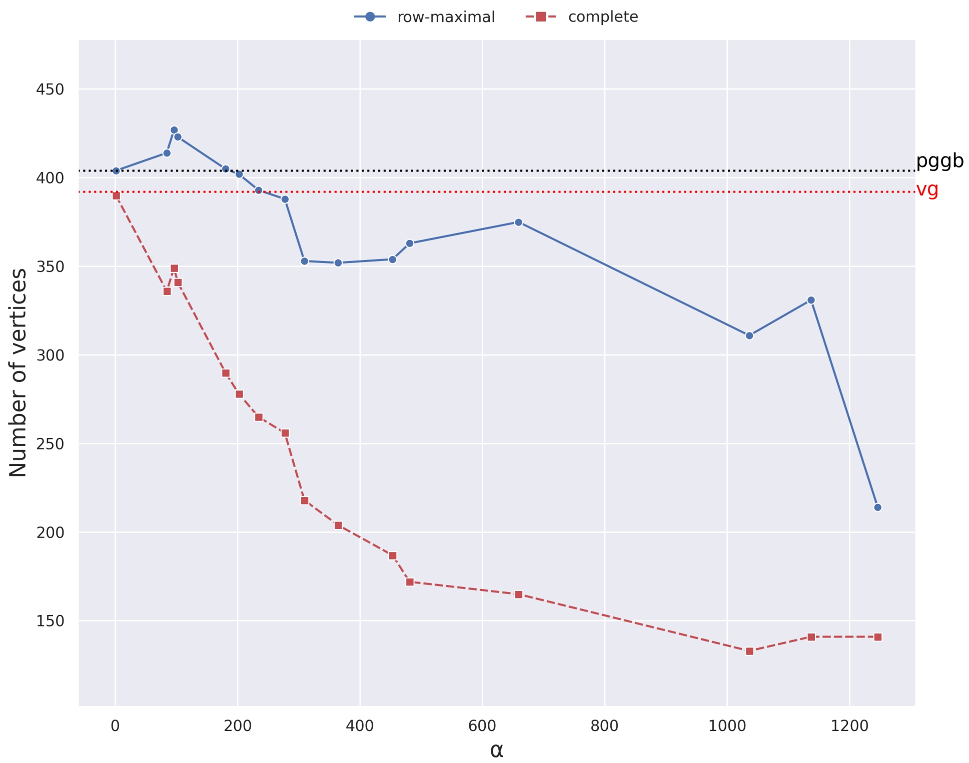


**Fig. 2**: **Number of vertices**. The plot is the results of running `pangeblocks`, `pggb`, and `vg` on the 20 SARS-CoV-2 genomes instance, using the row-maximal (blue dot) and the complete (red square) decomposition. The y-axis is the number of vertices of the variation graphs (after post-processing), and the x-axis has the objective functions. The black dotted and red dashed lines correspond to the number of nodes of the variation graph created with `pggb` and `vg`, respectively.

### 3 `pggb` and `vg`

**Table 2**: **Maximum RAM usage and User Time on SARS-CoV-2 MSAs** Results for 20, 50, and 100 sequences are shown for `pggb` and `vg`, both were run with their default parameters. Max RAM is in GB, and User Time is in seconds.

| | `pggb` | | `vg` | |
|---|---|---|---|---|
| Dataset | User Time | RAM | User Time | RAM |
| 20 | 62.83 | 0.998 | 0.34 | 0.026 |
| 50 | 184.05 | 1.246 | 1.58 | 0.062 |
| 100 | 599.71 | 1.808 | 7.65 | 0.135 |

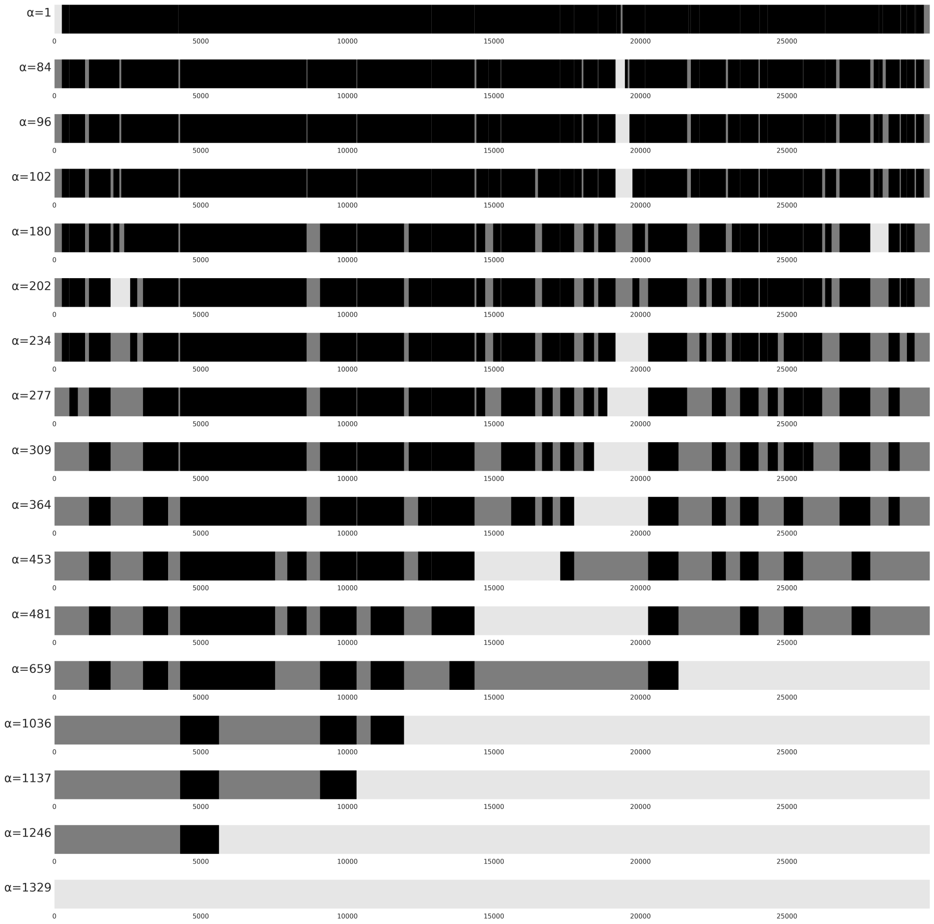


**Fig. 3**: A view of SARS-CoV-2 MSA with 20 sequences and 29879 columns for different values of $\alpha$ (see label to the left), from $\alpha = 1$ to $\alpha = 1329$ (from top to bottom). Forced vertical blocks (those with the number of columns less or equal to $\alpha$) are shown in black. SubMSAs in between vertical blocks (or at the beginning, or the end of the MSA) are in gray. The longest subMSA is shown in light gray. Values of $\alpha$ correspond to *breakpoints* shown in Figure 4

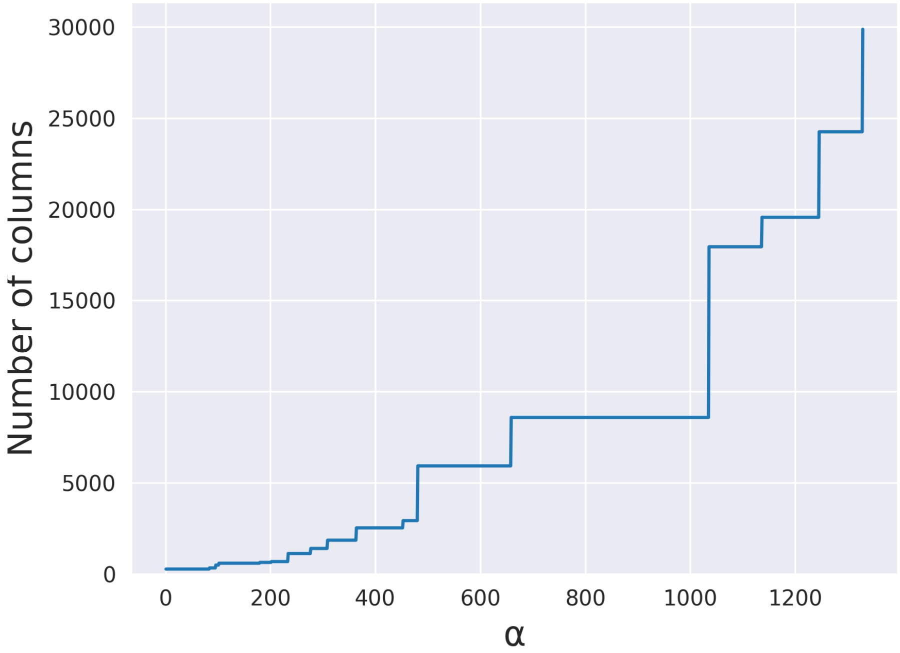


**Fig. 4**: Longest (columns) subMSA obtained for different values of $\alpha$ in the SARS-CoV-2 MSA with 20 sequences and 29879 columns. The vertical axis corresponds to the number of columns in the longest subMSA. The curve shows *breakpoints* of $\alpha$, *i.e.* where the number of columns in the longest subMSA increases.

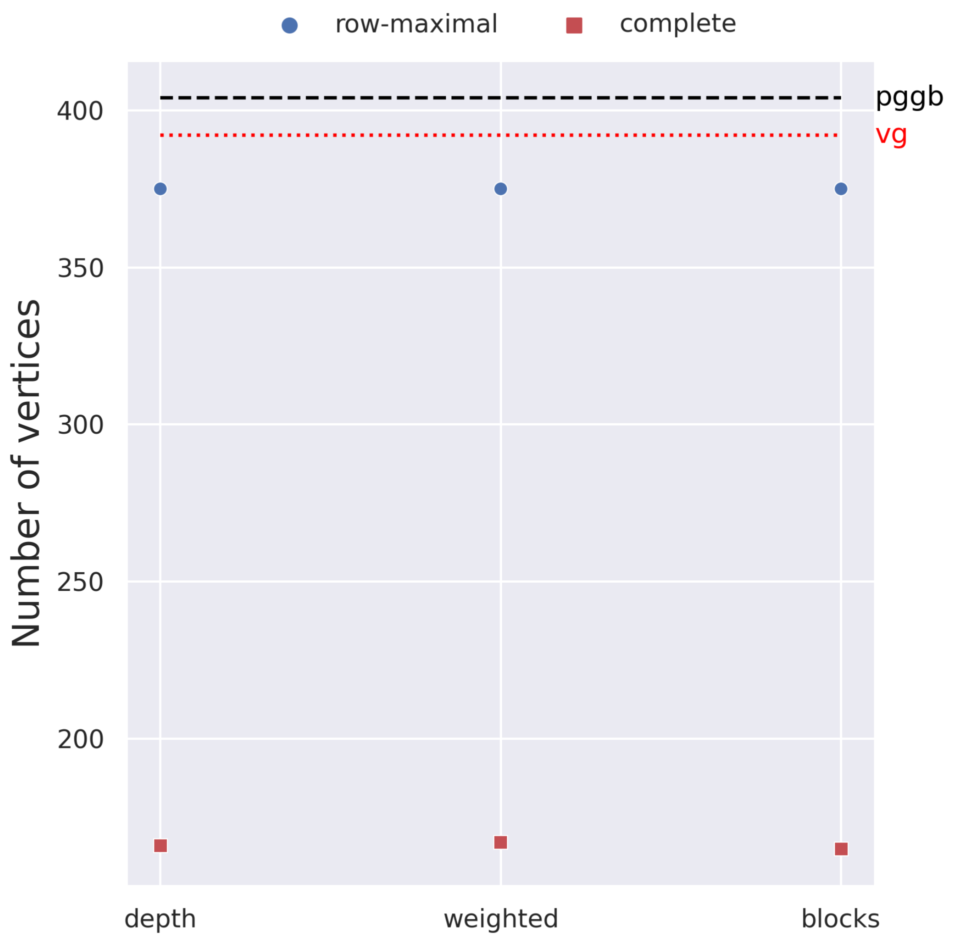


**Fig. 5**: **Number of vertices**. The plot is the results of running `pangeblocks`, `pggb`, and `vg` on the 20 SARS-CoV-2 genomes instance, using the row-maximal (blue dot) and the complete (red square) decomposition. The y-axis is the number of vertices of the variation graphs (after post-processing), and the x-axis has the objective functions. The black dotted and red dashed lines correspond to the number of nodes of the variation graph created with `pggb` and `vg`, respectively.

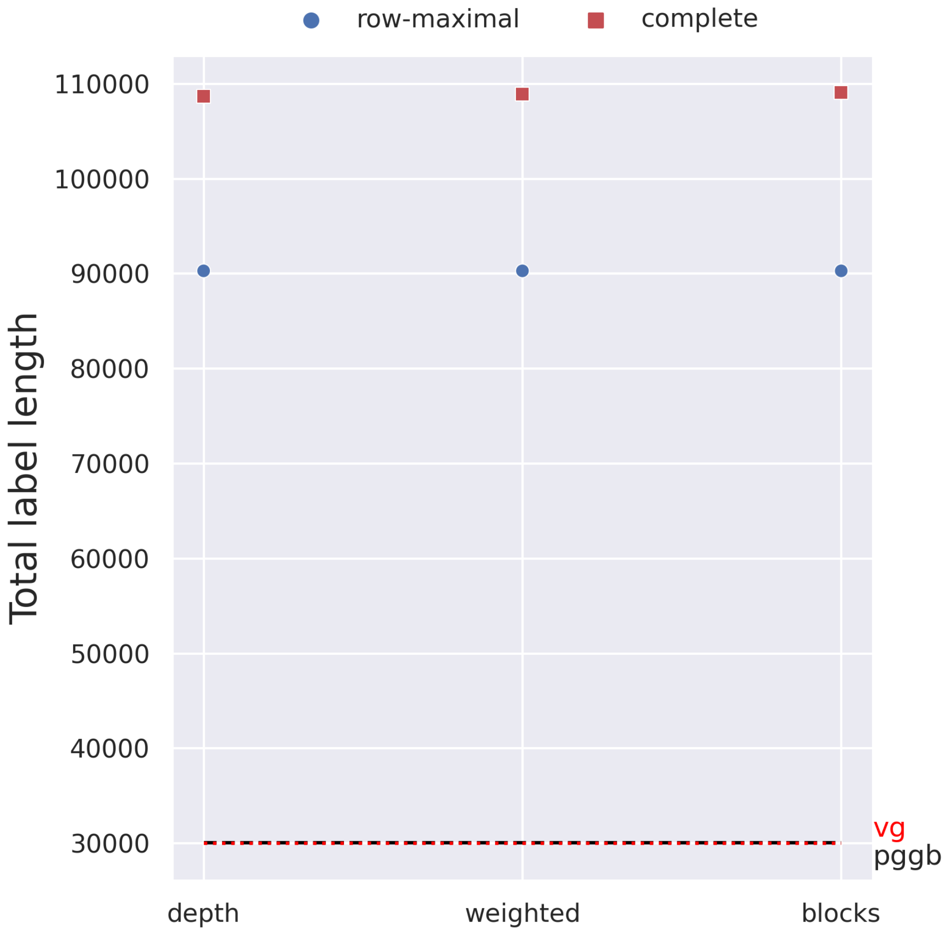


**Fig. 6**: **Length of the graph**. The plot is the results of running `pangeblocks`, `pggb`, and `vg` on the 20 SARS-CoV-2 genomes instance, using the row-maximal (blue dot) and the complete (red square) decomposition. The y-axis is the total number of characters of the graph labels (after post-processing), and the x-axis has the objective functions. The black dashed and red dotted lines correspond to the number of nodes of the variation graph created with `pggb` and `vg`, respectively.

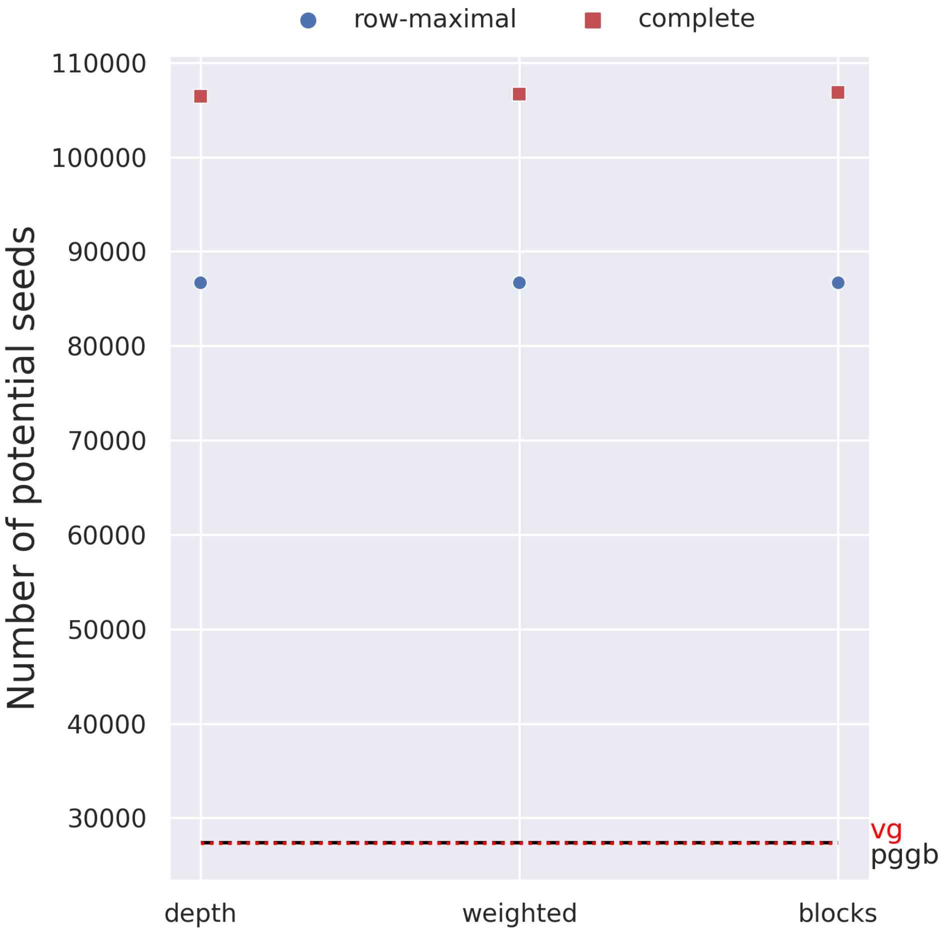


**Fig. 7**: **Number of potential seeds of length 20**. The plot is the result of running `pangeblocks`, `pggb`, and `vg` on the 20 SARS-CoV-2 genomes instance, using the row-maximal (blue dot) and the complete (red square) decomposition. The y-axis is the number of seeds (that is the 20-mers that are substring of at least a vertex label), the x axis is the $\alpha$ parameter. The black dashed and red dotted lines correspond to the results obtained by `pggb` and `vg`, respectively.

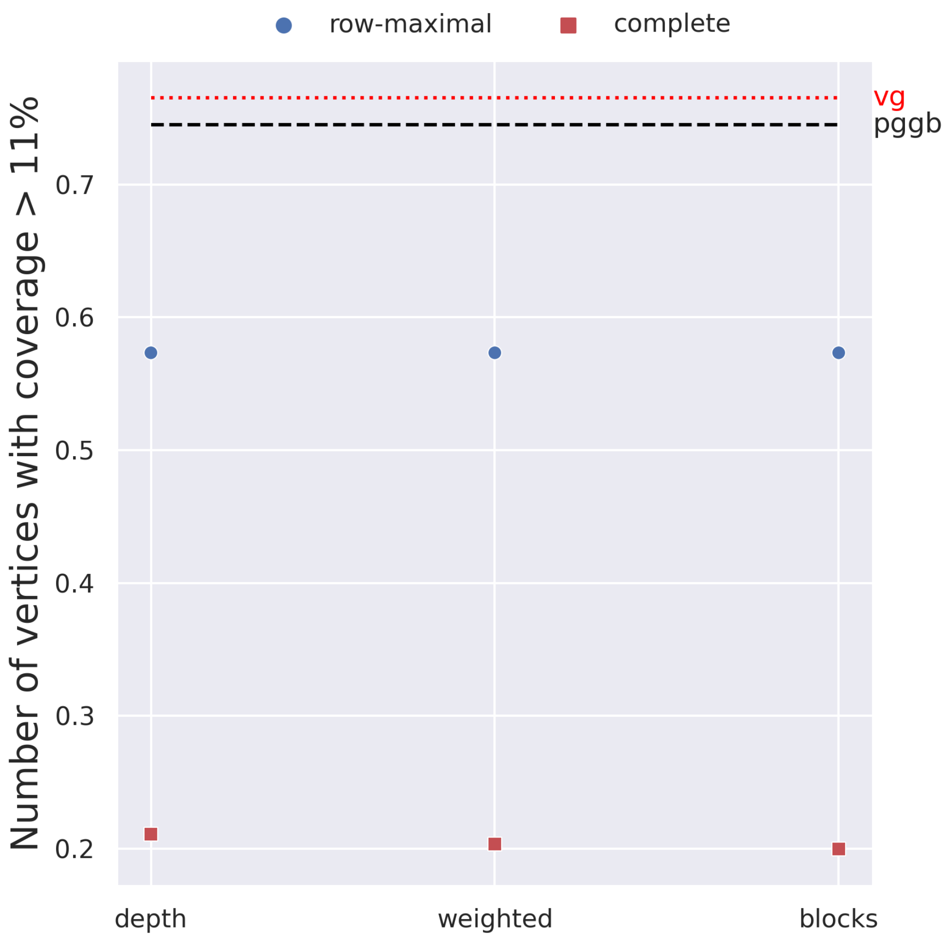


**Fig. 8**: **Number of vertices used by at least 11% of the genomes**. The plot is the result of running `pangeblocks`, `pggb`, and `vg` on the 20 SARS-CoV-2 genomes instance, using the row-maximal (blue dot) and the complete (red square) decomposition. The y-axis is the percentage of the vertices of the graph that traversed by more than 30% of the genome sequences. The black dashed and red dotted lines correspond to the number of nodes of the variation graph created with `pggb` and `vg`, respectively.

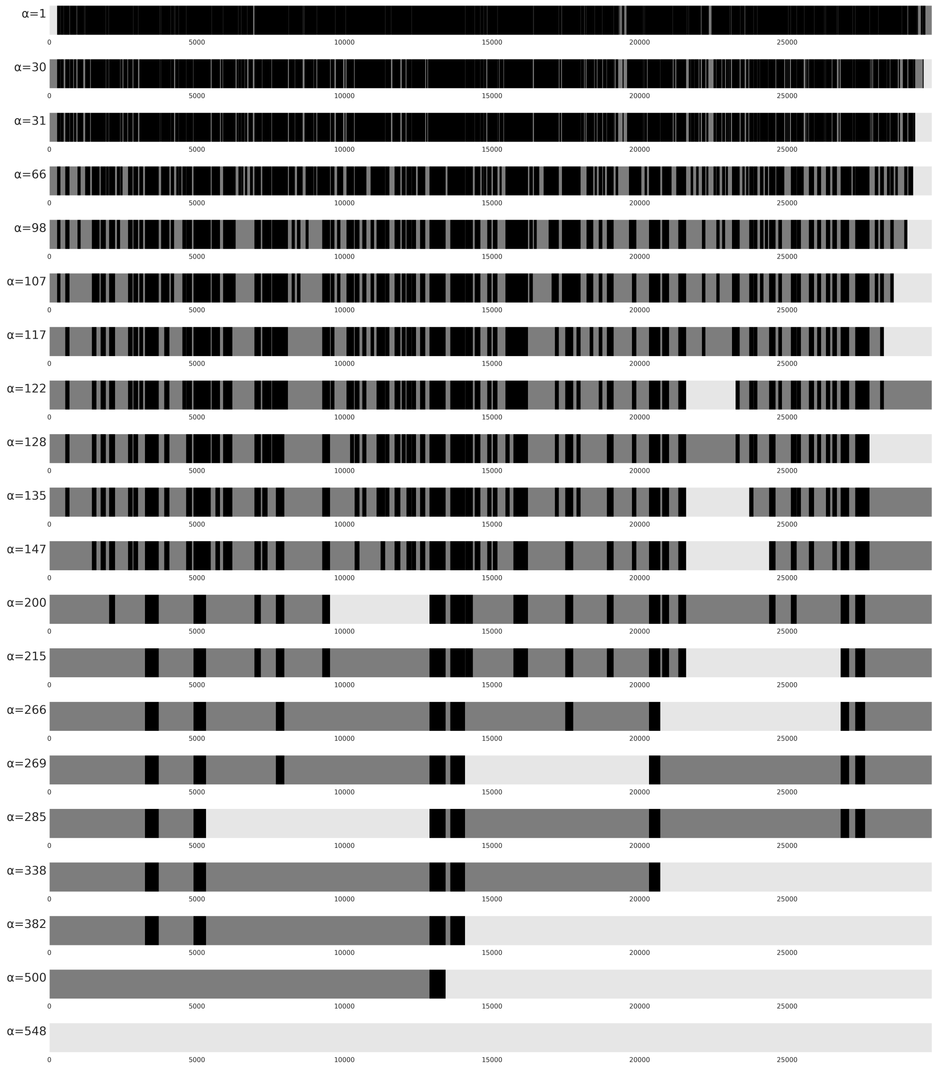


**Fig. 9**: A view of SARS-CoV-2 MSA with 100 sequences and 29904 columns for different values of $\alpha$ (see label to the left), from $\alpha = 1$ to $\alpha = 1329$ (from top to bottom). Forced vertical blocks (those with the number of columns less or equal to $\alpha$) are shown in black. SubMSAs in between vertical blocks (or at the beginning, or the end of the MSA) are in gray. The longest subMSA is shown in light gray. Values of $\alpha$ correspond to *breakpoints* shown in Figure 10

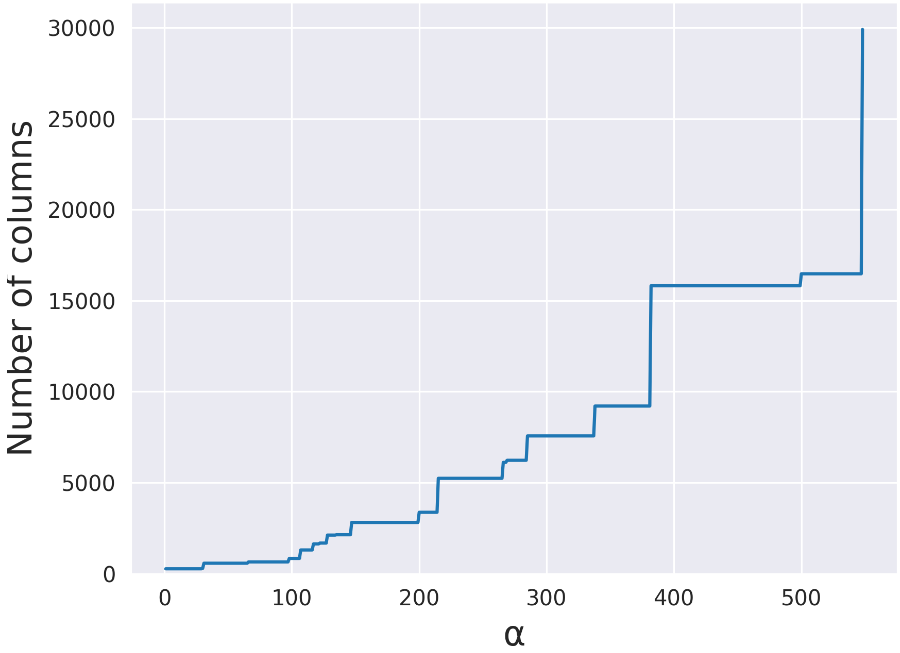


**Fig. 10**: Longest (columns) subMSA obtained for different values of $\alpha$ in the SARS-CoV-2 MSA with 100 sequences and 29904 columns. The vertical axis corresponds to the number of columns in the longest subMSA. The curve shows *breakpoints* of $\alpha$, *i.e.* where the number of columns in the longest subMSA increases.

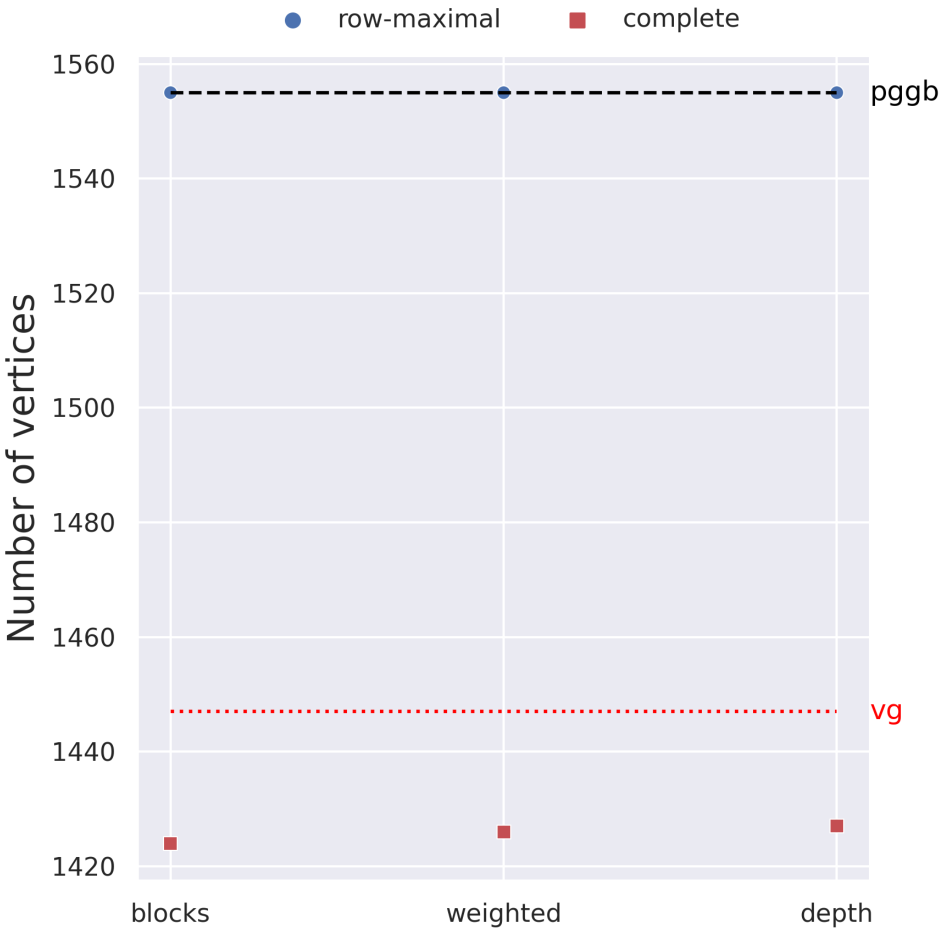


**Fig. 11**: **Number of vertices**. The plot is the results of running `pangeblocks`, `pggb`, and `vg` on the 100 SARS-CoV-2 genomes instance, using the row-maximal (blue dot) and the complete (red square) decomposition. The y-axis is the number of vertices of the variation graphs (after post-processing), and the x-axis has the objective functions. The black dotted and red dashed lines correspond to the number of nodes of the variation graph created with `pggb` and `vg`, respectively.

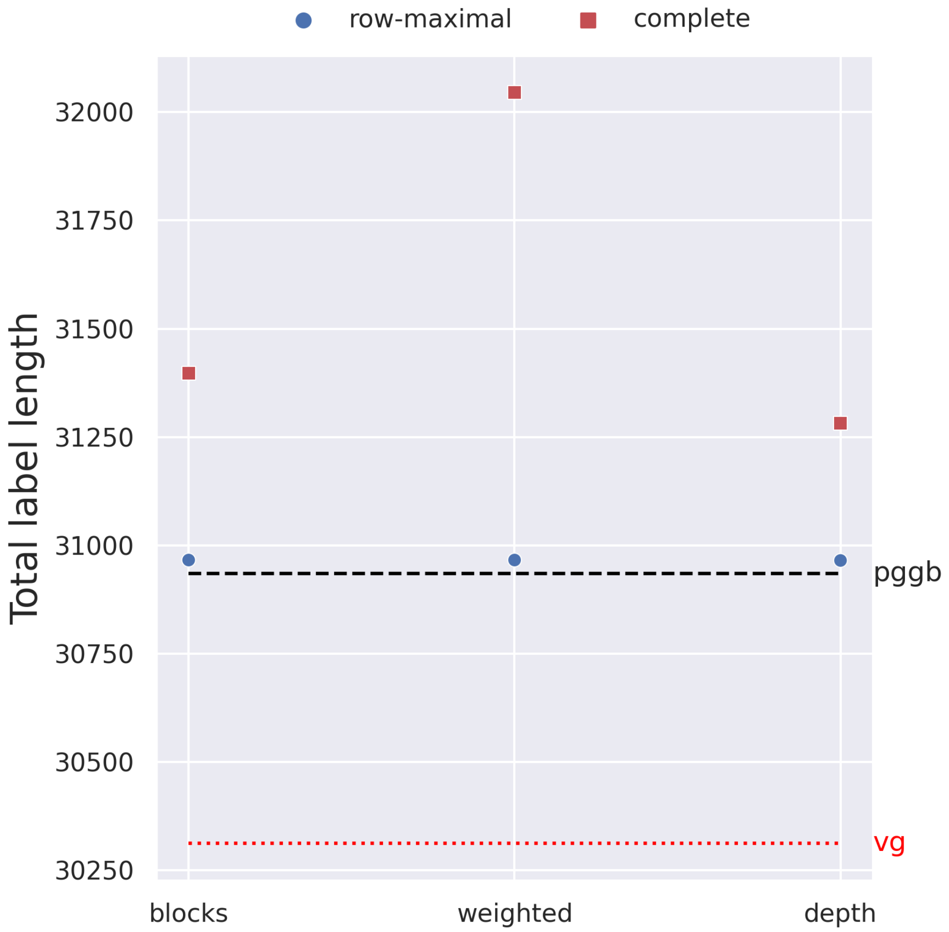


**Fig. 12**: **Length of the graph**. The plot is the results of running `pangeblocks`, `pggb`, and `vg` on the 100 SARS-CoV-2 genomes instance, using the row-maximal (blue dot) and the complete (red square) decomposition. The y-axis is the total number of characters of the graph labels (after post-processing), and the x-axis has the objective functions. The black dashed and red dotted lines correspond to the number of nodes of the variation graph created with `pggb` and `vg`, respectively.

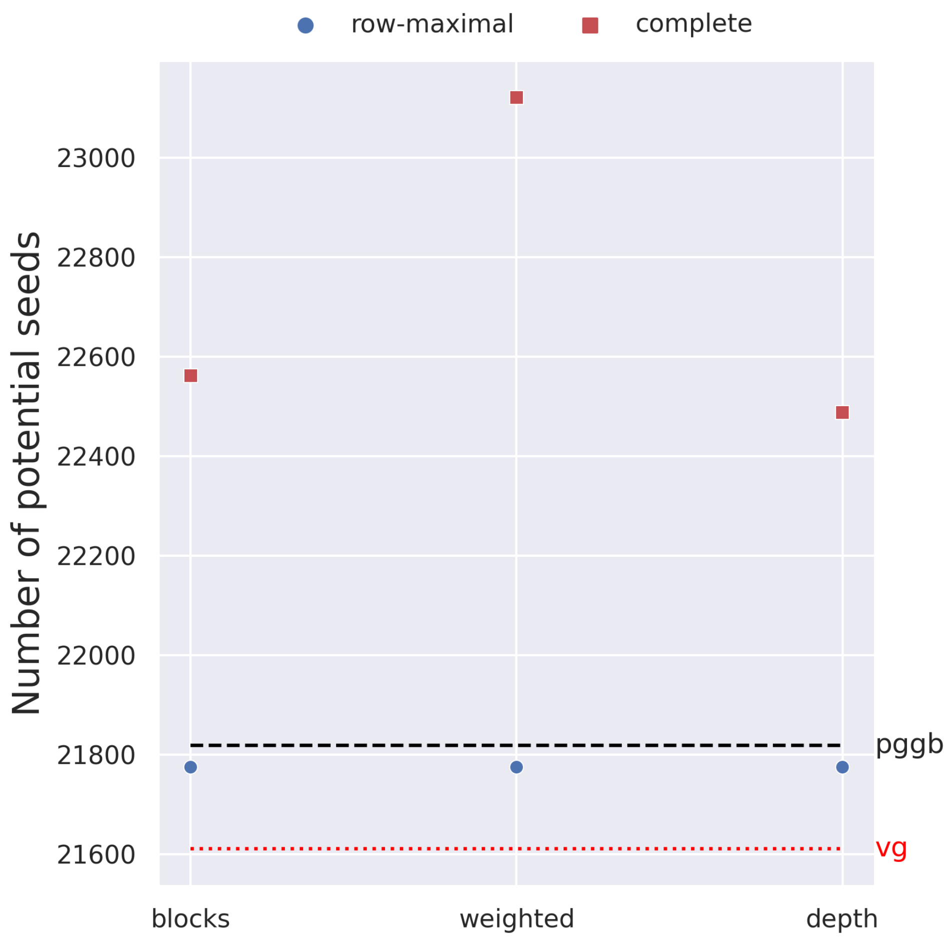


**Fig. 13**: **Number of potential seeds of length 20**. The plot is the result of running `pangeblocks`, `pggb`, and `vg` on the 100 SARS-CoV-2 genomes instance, using the row-maximal (blue dot) and the complete (red square) decomposition. The y-axis is the number of seeds (that is the 20-mers that are substring of at least a vertex label), the x axis is the $\alpha$ parameter. The black dashed and red dotted lines correspond to the results obtained by `pggb` and `vg`, respectively.

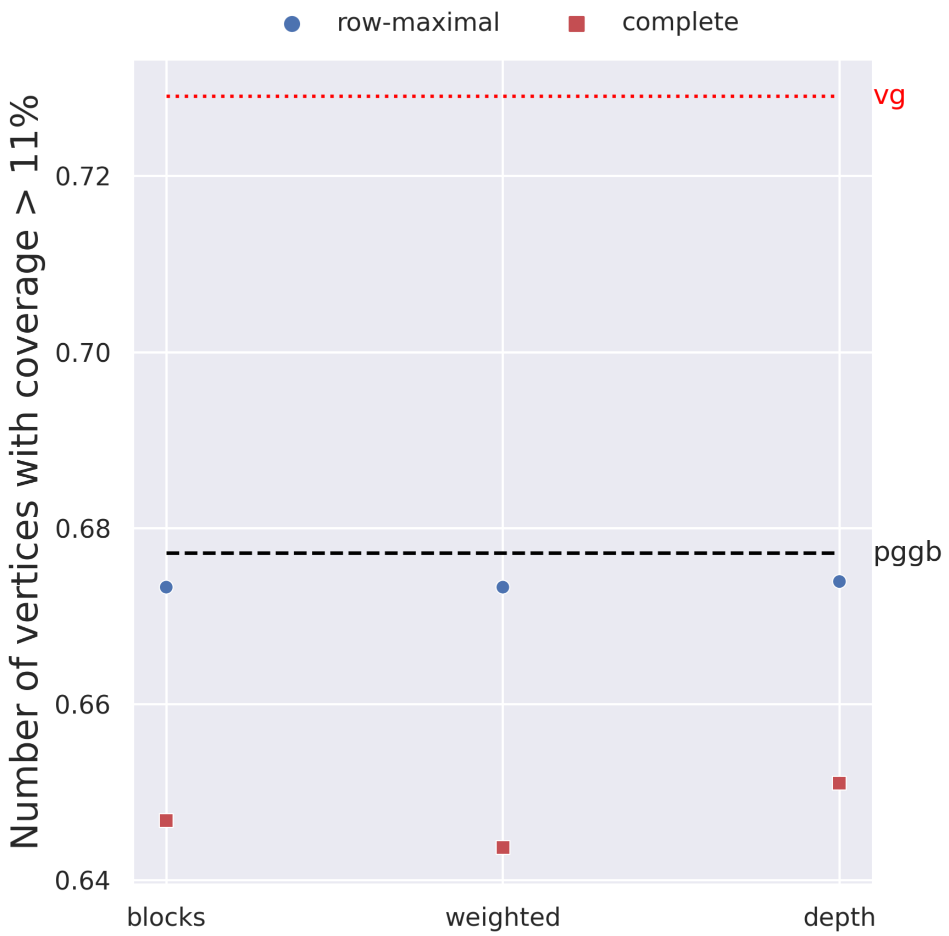


**Fig. 14**: **Number of vertices used by at least 11% of the genomes**. The plot is the result of running `pangeblocks`, `pggb`, and `vg` on the 100 SARS-CoV-2 genomes instance, using the row-maximal (blue dot) and the complete (red square) decomposition. The y-axis is the percentage of the vertices of the graph that traversed by more than 30% of the genome sequences. The black dashed and red dotted lines correspond to the number of nodes of the variation graph created with `pggb` and `vg`, respectively.
